# Large parallel deletions are associated with massive gene losses and virulence acquisition in *Meloidogyne incognita*

**DOI:** 10.64898/2026.09.04.749403

**Authors:** Ana Paula Zotta Mota, Hongli Yu, Maxime Multari, Rahim Hassanaly Goulamhoussen, Laetitia Perfus-Barbeoch, Sophie Valière, Kory Jason, Philippe Castagnone-Sereno, Pierre Abad, Cyril Van Ghelder, Etienne G J Danchin

## Abstract

The root-knot nematode *Meloidogyne incognita* is a major constraint to crop production worldwide. The resistance gene *Mi*-1.2 has been widely deployed in tomato varieties to control this parasite. However, virulent nematode isolates, overcoming the *Mi*-1.2-mediated resistance, have increasingly emerged in fields. Understanding the genetic mechanisms underlying resistance circumvention is therefore essential towards sustainable crop protection. From two geographically distinct *M. incognita* avirulent isolates, previous experimental evolution assays have allowed producing two virulent *M. incognita* isolates that escaped *Mi-*1.2 resistance in tomato. Here, we investigated the transcriptomic and genomic changes in the two virulent *M. incognita* isolates, as compared to their avirulent progenitor isolates controlled by the resistance. First, our transcriptomic analyses revealed hundreds of strongly downregulated genes, in common, in both virulent isolates. Most of these downregulated genes are clustered in two main genome regions. Long-read whole-genome sequencing of the four isolates revealed parallel structural variations in both virulent isolates. This included two large deletions spanning *ca.* 1.3 Mb and 0.6 Mb. The two big deletions encompass clusters of downregulated genes, suggesting they have in fact been lost in both virulent isolates. Short-read genome sequencing and PCR validation confirmed the two deletions in the two virulent isolates. The largest deletion includes predicted effector and orphan genes. The parallel large-scale structural variations identified here are likely to underlie the rapid adaptation of *M. incognita* to *Mi*-1.2-mediated resistance. These findings highlight genome structural plasticity as one of the potential key drivers of virulence acquisition in these asexually reproducing parasites.

## INTRODUCTION

Root-knot nematodes of the genus *Meloidogyne* are among the most destructive plant parasites in fields worldwide. Among them, *Meloidogyne incognita* is particularly damaging due to its global distribution, broad host range, and capacity to cause severe yield losses in vegetable production systems (J. T. Jones et al., 2013; Moura de Souza et al., 2026). *M. incognita* induces the formation of specialized feeding structures known as giant cells, leading to root galling, impaired nutrient uptake, and significant reductions in plant vigor and productivity (Abad et al., 2008). Plant genetic resistance represents one of the most effective and environmentally sustainable strategies to control root-knot nematodes. Since its introgression into cultivated tomatoes, the *Mi*-1.2 resistance gene has been widely deployed in commercial varieties and in many production systems. *Mi*-1.2 belongs to the NLR (nucleotide-binding domain leucine-rich repeat containing) family, which are pivotal actors of the effector-triggered immunity in plants. Therefore, *Mi*-1.2, directly or indirectly detects parasites and triggers an hypersensitive response in the tomato, thus preventing parasites from developing (Milligan et al., 1998). However, like many single dominant resistance genes, *Mi*-1.2 exerts strong selective pressure on parasite populations. Experimental evolution and field evidence have both shown that virulent *M. incognita* isolates capable of overcoming *Mi*-1.2 can emerge after repeated exposure to resistant cultivars (Barbary et al., 2015; Castagnone-Sereno et al., 1994). The escape from plant resistance threatens the long-term durability of this pivotal control strategy and underscores the need to understand the genetic mechanisms of virulence acquisition that enable nematodes to evade host immunity.

*M. incognita* exhibits a complex triploid genome structure (Dai et al., 2023; Mota et al., 2024), and reproduces exclusively via parthenogenesis (Koutsovoulos et al., 2020; Mota et al., 2024). Despite its clonal reproduction and absence of outcrossing, the species displays remarkable adaptive potential. Genome plasticity, including single nucleotide polymorphisms (SNPs) (Koutsovoulos et al., 2020; Szitenberg et al., 2017), structural variation (Castagnone-Sereno et al., 2019), and transposable element (TE) activity (Kozlowski et al., 2020), may contribute to its adaptability, together with epigenetic factors (Dai et al., 2026; Hassanaly-Goulamhoussen et al., 2021). A previous study, using comparative genomic hybridization (CGH) array and qPCR, based on two pairs of experimentally developed virulent / avirulent near-isogenic lines (NILs), confirmed 18 parallel gene copy losses shared by the two virulent lines, despite distant geographic origins (Castagnone-Sereno et al., 2019). However, these CGH array studies were built from an initial incomplete version of the genome assembly, in which several gene families and gene copies were missing (Abad et al., 2008; Blanc-Mathieu, Perfus-Barbeoch, Aury, Da Rocha, et al., 2017). Furthermore, no systematic analysis of gene expression differences at the whole-genome scale between virulent and avirulent isolates were conducted.

Therefore, in this study, we used the same two virulent / avirulent pairs of NILs, to comprehensively investigate the transcriptomic and genomic changes associated with virulence acquisition by *M. incognita*. We combined transcriptome and whole-genome sequencing to infer parallel changes associated with resistance circumvention using a highly complete long-read-based *M. incognita* reference genome (Mota et al., 2024). We identified 707 differentially expressed genes (DEGs) in the virulent isolates, as compared to the avirulent ones, of which 639 were downregulated. Genome analyses allowed identification of two large megabase-scale deletions encompassing 451 downregulated genes that are shared by the two virulent isolates. The DEGs in the deletions encoded several putative effectors that may be involved in the *Mi*-1.2 mediated recognition and plant defense. Their deletion might allow them escape from *Mi*-mediated host resistance.

## MATERIAL AND METHODS

### Avirulent and virulent near-isogenic nematode isolates

Two avirulent *M. incognita* isolates controlled by the tomato *Mi*-1.2 resistance gene (Vos† et al., 1998) were originally sampled in the field in geographically distinct locations, Kursk, in Russia, and Morelos, in Mexico. In each isolate, one single female was characterized to confirm correct species identification and its egg mass was then dissected and reared on a susceptible tomato plant, lacking *Mi-*1.2, under greenhouse conditions. The second-stage juveniles that hatched from the egg mass were then used as a starting clonal line. From each avirulent line, a near-isogenic line (NIL) was selected for virulence by successive re-inoculation on the *Mi*-resistant tomato cultivar Piersol. Nematodes on resistant plants eventually became able to circumvent the plant immunity and establish feeding sites. In parallel, the rest of the original avirulent isolate, was maintained on the susceptible tomato cultivar Saint Pierre. These two newly derived virulent isolates (Morelos virulent, MV; Kursk virulent, KV), together with the two original near-isogenic avirulent isolates (Morelos avirulent, MA; Kursk avirulent, KA) were successively maintained on the corresponding tomato cultivars, by planting new tomato plants every two months. Detailed experiment procedures are described in (Castagnone-Sereno et al., 2019).

### Transcriptome sequencing and data processing

Total RNA was isolated using the Trizol extraction method for each of the NILs (MA, MV, KA and KV). RNA-seq libraries were prepared by Genome Quebec and sequenced using an Illumina NovaSeq 6000 with 2 x 100 bp paired-end reads.

The quality of the sequenced raw reads was assessed using FastQC (Andrews, 2010) and summarized with MultiQC (Ewels et al., 2016). Reads with a mean Q score < 28 and a length < 75 bp were removed (**Supplementary Table S1**). Five bp from the beginning and 3 bp from the end were trimmed as the quality tends to be lower at these ends. One replicate for population KA was later identified as outlier based on principal component analysis and was not included in the final analysis. We mapped the clean reads of each library against the long-read *M. incognita* reference genome (Mota et al., 2024), using STAR (Dobin et al., 2013) with default parameters. We further counted the mRNA feature and mapped reads using featureCounts (-t mRNA) from the mapping files and grouped them by ID attribute (-g ID), producing one featureCounts table per sample. We then merged the featureCounts tables to obtain one table spanning all the samples. We used DESeq2 (Love et al., 2014) to detect differentially expressed genes (DEGs). As the objective of this study was to detect differences between avirulent and virulent isolates, data from the two avirulent isolates (KA, MA) were pooled and compared to pooled data of the two virulent ones (MV, KV) regardless of their geographic origins (design = ∼genotype in DESeq2). We used the Independent Hypothesis Weighting (IHW) model for p-value correction and considered as DEGs all the genes with adjusted p-value ≤ 0.05 and |log_2_ (fold change)| ≥ 1. DEGs between virulent and avirulent populations, and distributions along contigs were plotted using ggplot2 v. 3.3.5 (Wickham, 2009). Functional annotation was based on the InterproScan analysis from the reference genome (Mota et al., 2024).

### Whole-genome sequencing and data processing

For each of the four isolates, genomic DNA was extracted from purified eggs. High molecular weight DNA for Oxford Nanopore Technologies (ONT) long-read sequencing was extracted using the Epicentre MasterPure Complete DNA and RNA Purification Kit (Lucigen, Biosearch Technologies, Middleton, WI, USA) according to the manufacturer’s instructions, with vortexing steps omitted to preserve DNA integrity. For the MA population, we directly used the long reads generated to produce the *M. incognita* reference genome (Mota et al., 2024), as they are from the exact same Morelos avirulent (MA) population. Library preparation and whole-genome nanopore sequencing were performed at the UCAGenomiX platform using a PromethION Oxford Nanopore Technology (ONT) instrument. ONT libraries were generated using the SQK-LSK109 ligation kit as per Oxford Nanopore instructions. The libraries were loaded into R9 flow cells.

Genomic DNA for AVITI short-read sequencing and PCR molecular validation was extracted using an in-house CTAB protocol as detailed below. Ground eggs were resuspended in extraction buffer (2% CTAB, 1.4 M NaCl, 20 mM EDTA, 100 mM Tris-HCl, pH 8.0, and 0.2% β-mercaptoethanol) and incubated at 65°C for 30 min. Samples were then treated with RNase and then Chloroform: isoamyl alcohol (24: 1), followed by centrifugation at 10,000 x *g* for 10 min. Supernatants were collected, isopropanol was added, and the samples were stored at −20°C overnight. Samples were then centrifuged and the resulting pellets were washed twice with ethanol 70% before being resuspended in pure sterile water. The concentration and purity of the extracted DNA were assessed using a Qubit fluorometer and a NanoDrop spectrophotometer.

High-coverage AVITI whole-genome sequencing was performed at the GeT-PlaGe core facility, INRAE Toulouse. Sequencing libraries were prepared using the Illumina TruSeq Nano DNA HT Library Prep Kit. Briefly, DNA molecules were fragmented by sonication using the PIXUL® Multi-Sample Sonicator (ActiveMotif, 53130) for a target insert size of 550 bp, followed by adapter ligation. Seven cycles of PCR were applied to amplify the libraries. Library quality was assessed using an Advanced Analytical Fragment Analyzer (Agilent) and libraries were quantified using Qubit 3.0 (Thermo Fisher Scientific). Sequencing was performed on an Element Biosciences AVITI using a paired-end read length of 2 x 150 pb with the High Output Cloudbreak Freestyle 300 cycles.

Length and quality of ONT reads were first checked using cONTent, a tool available at https://github.com/DjampaKozlowski/cONTent, and filtered for at least 1 kb length (-m 1000) and a min. quality score of Q12 (-Q 12). This allowed having a minimum genome coverage of 27x (mean 66x) for all the samples. Summary of read counts and length are provided in **Supplementary Table S2.** The processed reads were then mapped to the reference genome using Minimap2 v. 2.24 (Li, 2018) with the ONT-specific preset (-ax map-ont) and Samtools (Li & Durbin, 2009), filtering for mapping quality (-q 12). The final mapped read depth per base was calculated using Samtools (-depth) with default settings, which was then summarized based on 1 kb sliding window and plotted using ggplot2.

AVITI reads were trimmed and filtered using fastp v. 0.23.4 (S. Chen et al., 2018). Low-quality bases from both the 5′ and 3′ ends were trimmed, reads containing more than 28% low-quality bases were discarded, and reads shorter than 120 bp after trimming were removed (-5 -3 -M 28 -l 120). Summary of read counts and length are provided in **Supplementary Table S2.** The cleaned reads were then mapped to the reference genome using the BWA-MEM2 v. 2.2.1 (Li & Durbin, 2009) mem algorithm and Samtools, filtering for mapping quality (-q 20). The final mapped read depth was calculated and plotted with the same methods used for the ONT data.

### PCR validation of deletions and species identification

PCR validation assay was used to confirm the observed deletions in virulent populations. Primers were designed using Primer3Web v. 4.1.0 with default settings (**Supplementary Table S3**). PCR was performed using MyTaq kit (Meridian Bioscience, Bioline, London, UK), following supplier’s instructions. PCR conditions were as follows: 94°C for 2 min, followed by 35 cycles of 94°C for 30 s, 59°C for 30 s, and 72°C for 45 s, with a final extension step at 72°C for 5 min. Species identification was confirmed using the *M. incognita* SCAR marker Inc-K-14 following the instructions described in (Randig et al., 2002).

### Functional annotation, structural prediction and GO enrichment

The functional annotation is based on the InterProScan 5 analysis (P. Jones et al., 2014) from the long-read reference genome (Mota et al., 2024), and is available in (https://entrepot.recherche.data.gouv.fr/dataverse/Ma-Mv-Ka-Kv/). Gene Ontology (GO) terms enrichment analysis was conducted using the R package GOfuncR (https://doi.org/doi:10.18129/B9.bioc.GOfuncR). The MAP-1 protein sequence was used as a query in BlastP analysis (e-value< 1e-50) to retrieve homologs from reference predicted proteomes (https://entrepot.recherche.data.gouv.fr/dataverse/gene-preds and https://entrepot.recherche.data.gouv.fr/dataverse/Ment-HiFI-E1834) (Mota et al., 2024; Poullet et al., 2025). InterProScan (P. Jones et al., 2014) results were used to assess presence / absence of domain / motif in the obtained sequences to only retain sequences containing a signal peptide, a disorder proline-rich region and a Barwin-like endoglucanases domain. A MAP-1 phylogenetic tree was produced using MEGA X (Kumar et al., 2018) with a multiple sequence alignment performed with Muscle (Edgar, 2022). The tree was produced with the Maximum Likelihood method and JTT matrix-based model. The percentage of trees (100 bootstraps) in which the associated taxa clustered together is shown next to the branches. Initial tree(s) for the heuristic search were obtained by applying Neighbor-Joining in the BioNJ algorithm to a matrix of pairwise distances estimated using the JTT model, and then selecting the topology with superior log likelihood value. This analysis involved 11 protein sequences. There were a total of 523 positions in the final alignments. Secondary and 3D structures of MAP-1 were predicted using JPred 4 (Drozdetskiy et al., 2015) and AlphaFold 3 (Abramson et al., 2024).

## RESULTS

### A massive number of down-regulated genes in both virulent isolates

Starting from two originally avirulent *M. incognita* isolates (Morelos avirulent, MA; Kursk avirulent, KA), two near-isogenic virulent isolates (Morelos virulent, MV; Kursk virulent, KV) were previously obtained via experimental evolution (Castagnone-Sereno et al., 2019). These virulent isolates are able to develop on tomatoes carrying the widely-used *Mi-*1.2 resistance gene. Infection symptoms observed on *Mi-*1.2*-*carrying tomatoes were similar to those regularly noted on susceptible tomato plants (**Supplementary Figure S1**). We sequenced the transcriptomes of the four lines (MA, MV, KA, KV) in triplicates. One library of KA was identified as an outlier based on principal component analysis and was not included in the final analysis, resulting in 11 libraries. An average of 35.8 million (range from 28.2 in a KV library to 40.2 in a MV library) clean read pairs were obtained per library, from the original 36.3 million (range from 28.6 in a KV library to 40.7 in a MV library) 2 x 101 bp raw read pairs. Summary of data statistics is included in **Supplementary Table S1**. We conducted a differential gene expression analysis, comparing libraries from the virulent isolates to those from the avirulent isolates. A total of 707 differentially expressed genes (DEGs) were identified in virulent isolates as compared to avirulent ones, of which 639 were downregulated and 68 upregulated (**Figure 1a; Supplementary Table S4**). These DEGs were distributed across 113 contigs in the reference genome, with five contigs containing nearly half of the DEGs (352 out of 707). Visualization of DEG distribution along the five contigs identified high densities of DEGs in the beginning of contig 2 and near the end of contig 25 (**Figure 1b**). Contigs 2 and 25 also comprise the largest number of DEGs (286 and 29, respectively), accounting for 40.5% and 4.1% of the total identified, respectively, and notably, all the DEGs in these two contigs are downregulated in both virulent isolates.

**Figure 1.**
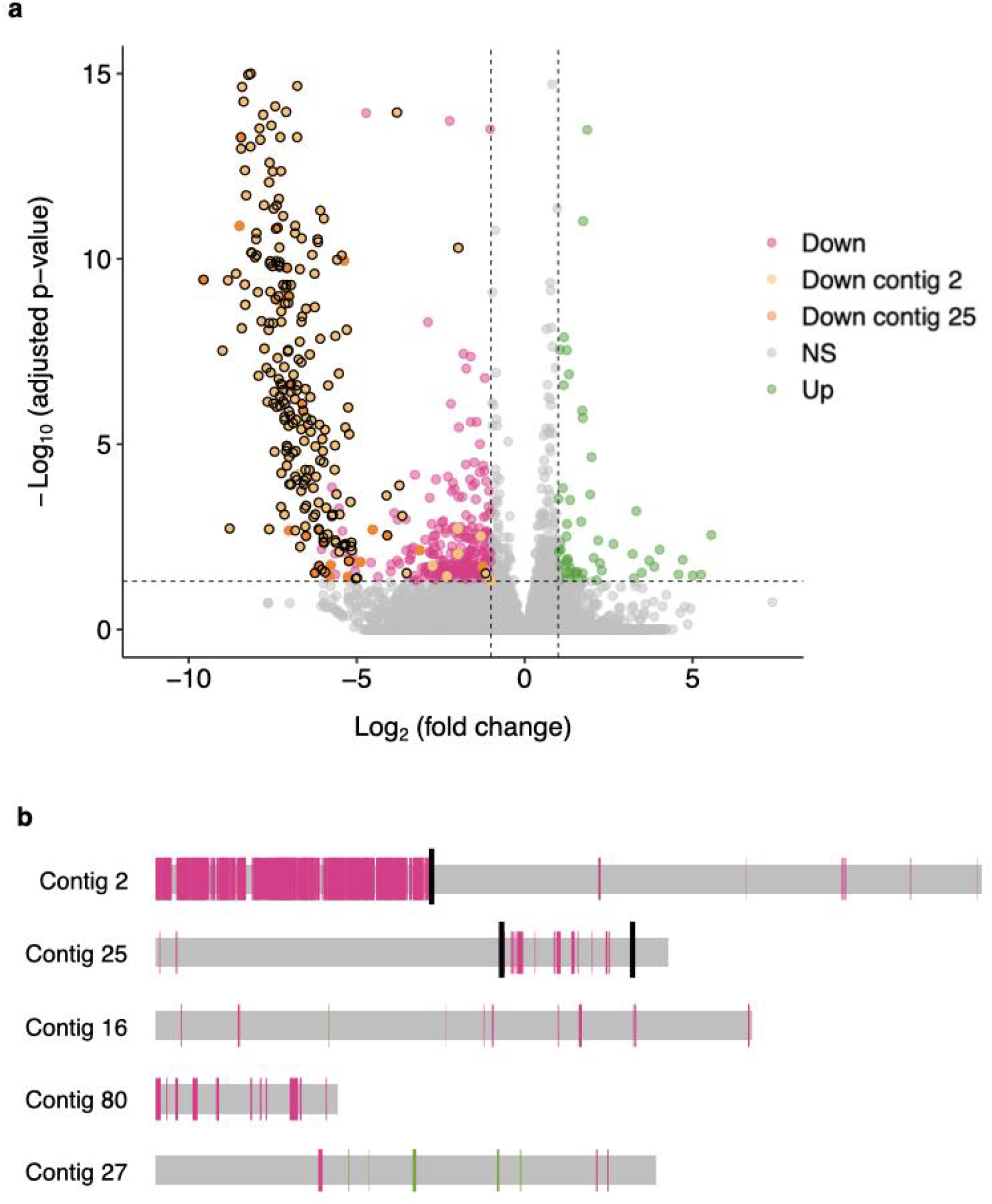
Differentially expressed genes (DEGs) in virulent populations and their distribution in the *M. incognita* genome. (**a**) Volcano plot showing DEGs in virulent versus avirulent isolates. Genes with adjusted p-value ≤ 0.05 and |log_2_ (fold change)| ≥ 1 are marked with light orange (downregulated in contig 2), dark orange (downregulated in contig 25), pink (downregulated in other contigs), and green (upregulated), while those not significantly (NS) differentially expressed are marked as grey. Points outlined in black indicate genes within the identified large deletions in contig 2 and contig 25 (next section). No genes present on contig 2 and contig 25 were upregulated. (**b**) Distribution of DEGs along the five contigs containing the largest numbers of DEGs. Pink, and green bars represent DEGs that are downregulated, and upregulated, respectively. Grey horizontal bars indicate contigs and are proportional to their actual lengths. Contigs are ranked from top to bottom by descending number of DEGs. High densities of DEGs were found at the beginning of contig 2 and near the end of contig 25, as delineated by the black lines.

Gene Ontology (GO) enrichment analysis on DEGs revealed that various ‘molecular function’ GO terms related to peptidase activities were significantly enriched in downregulated genes (**Supplementary Data Table S5**). At the ‘biological process’ level, GO terms related to histone methylation and endosome regulation were enriched in downregulated genes, together with the ‘cellular component’ term microtubule. On the other hand, the ‘biological process’ GO term ‘protein catabolic process’ was the only one enriched in upregulated genes, while no GO terms in the categories ‘molecular function’ or ‘cellular component’ were significantly enriched. Based on the InterProScan annotation, we categorized the proteins that contained signal peptides but no transmembrane domains (SP-noTM) as predicted secreted proteins. The SP-noTM dataset is therefore presumably enriched in effectors secreted by the nematode. A total of 10.2 % (65 out of 639) of the downregulated genes belong to this dataset, slightly higher than the whole predicted proteome (8.1%) (even though not statistically enriched; *□*2; P = 0.063), whereas only 2.6% are found in the upregulated genes. Besides, some downregulated genes are homologs of experimentally confirmed nematode effectors. For instance, the Minc_v4_shac_contig_108g0488901 gene (Log_2_ (fold change) = -2.1; Padj = 1.7 x10^-3^) is an homolog (86% identity at the protein level) of PEL-1 (AAS88579), a pectate lyase expressed in pre-parasitic and parasitic *M. incognita* J2 sub-ventral glands. While Minc_v4_shac_contig_80g0440501 (Log_2_ (fold change) = -3.4; Padj = 1.3 x10^-2^ is an homolog (98% identity) of the pioneer putative effectors msp6 (AF531165), msp13 (AY134432) and msp23 (AY134442) expressed in dorsal glands (Huang et al., 2003). Similarly, Minc_v4_shac_contig_103g0480431 (log_2_ (fold change) = -2.1; Padj = 1.1 x10^-4^) is an homolog of Minc01696 (88% identity), a putative secreted protein kinase expressed in sub-ventral glands of *M. incognita* (Rutter et al., 2014). Candidate effectors located on contigs 2 and 25 are presented in a separate section in the manuscript.

### Large parallel genome deletions identified in virulent populations

As high densities of downregulated genes in virulent isolates compared to avirulent ones were observed in certain genomic regions (e.g., the beginning of contig 2 and near the end of contig 25), we further investigated whether some structural variations were present in these genome regions. We performed whole-genome sequencing for the four isolates using both long- and short-read sequencing approaches. After filtering, ONT sequencing generated on average 1.5 million (range from 0.3 in KA to 2.6 in MA) clean reads per isolate, with an average length of 8.9 kb, and AVITI sequencing generated on average 136 million (range from 104.8 in MA to 161.1 in MV) read pairs of 150 bp per isolate. Summary of data statistics is included in **Supplementary Table S2**. We mapped all these whole-genome sequencing reads to the MA reference genome (Mota et al., 2024), and first plotted the read depth distribution for contigs 2 and 25. Results from both ONT and AVITI data indicate that regions of *ca.* 1.3 Mb (1 bp to 1.2655 Mb for KV and 1 bp to 1.3035 Mb in MV) at the beginning of contig 2 and *ca.* 0.6 Mb (1.5855–2.1855 Mb) near the end of contig 25 have very low to no coverage by reads from the two virulent isolates (KV, MV), while relatively consistent coverage is observed in the two avirulent ones (KA, MA) **(Figure 2)**. Likewise, we plotted the read depth distribution for the 100 largest contigs (**Supplementary Figure S2**). Similar situations of regions with little to no read coverage in the two virulent isolates despite consistent coverage in the two avirulent ones were also observed in some other contigs (e.g., contig 80), suggesting other deletions associated with virulent populations.

**Figure 2.**
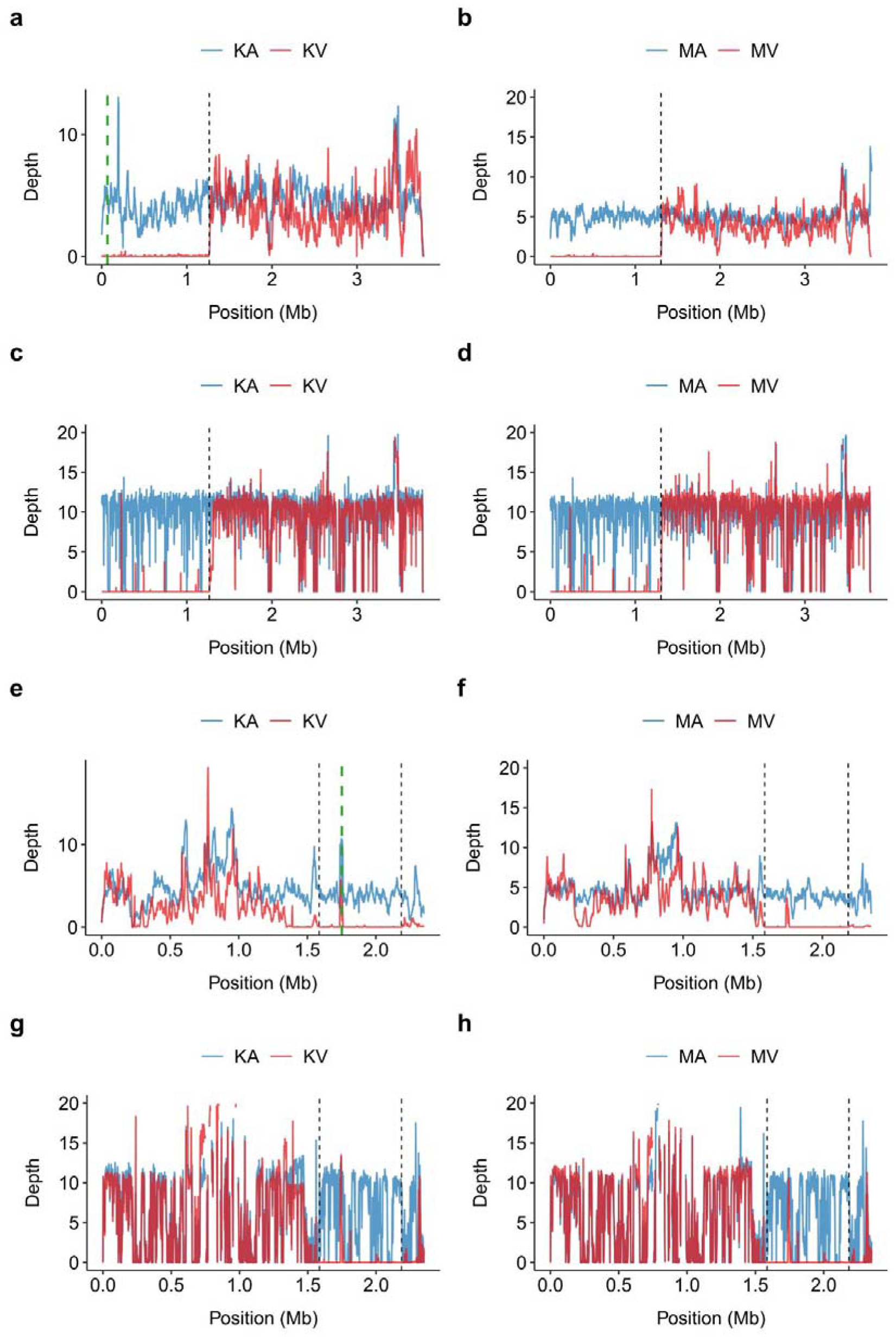
Mapping depth of whole-genome sequencing reads on contig 2 (**a**-**d**) and 25 (**e**-**h**) for avirulent (KA, MA; blue) and virulent (KV, MV; red) isolates showing deleted regions in contig 2 (from beginning to the black dashed line; *ca.* 1 bp to 1.2655 Mb in KV and 1 bp to 1.3035 Mb in MV) and contig 25 (between the black dashed lines; 1.5855–2.1855 Mb) in virulent isolates. **a, b, e,** and **f** show read depths from Oxford Nanopore long reads, while **c, d, g** and **h** show read depths from AVITI short reads. Depth is the averaged mapping depth of reads per kb window and per Gb bam file size. Green dashed lines in panel **a** and panel **e** indicate locations of primers used in PCR validation of the deletions.

As the two identified large deleted regions in contig 2 and contig 25 combined account for *ca.* 1% of the total length of the *M. incognita* reference genome and contain the highest densities of DEGs, we focused on these two regions in the following analysis. Notably, 340 protein-coding genes are present in the *ca.* 1.3 Mb deletion in contig 2, and were probably lost in parallel in the two virulent isolates. Similarly, 111 genes were present in the *ca.* 0.6 Mb deletion in contig 25 and were probably lost in both virulent isolates. Using a SCAR marker, we confirmed the four isolates are indeed all *M. incognita*. Then, a PCR experiment using primers targeting the two deleted regions confirmed their absence in the virulent isolates while they were amplified in the avirulent ones. An independent neutral control designed in contig 1 showed consistent amplification in all four isolates **(Figure 3)**. Thus, these results confirmed the parallel loss of genomic material in the two virulent isolates.

**Figure 3.**
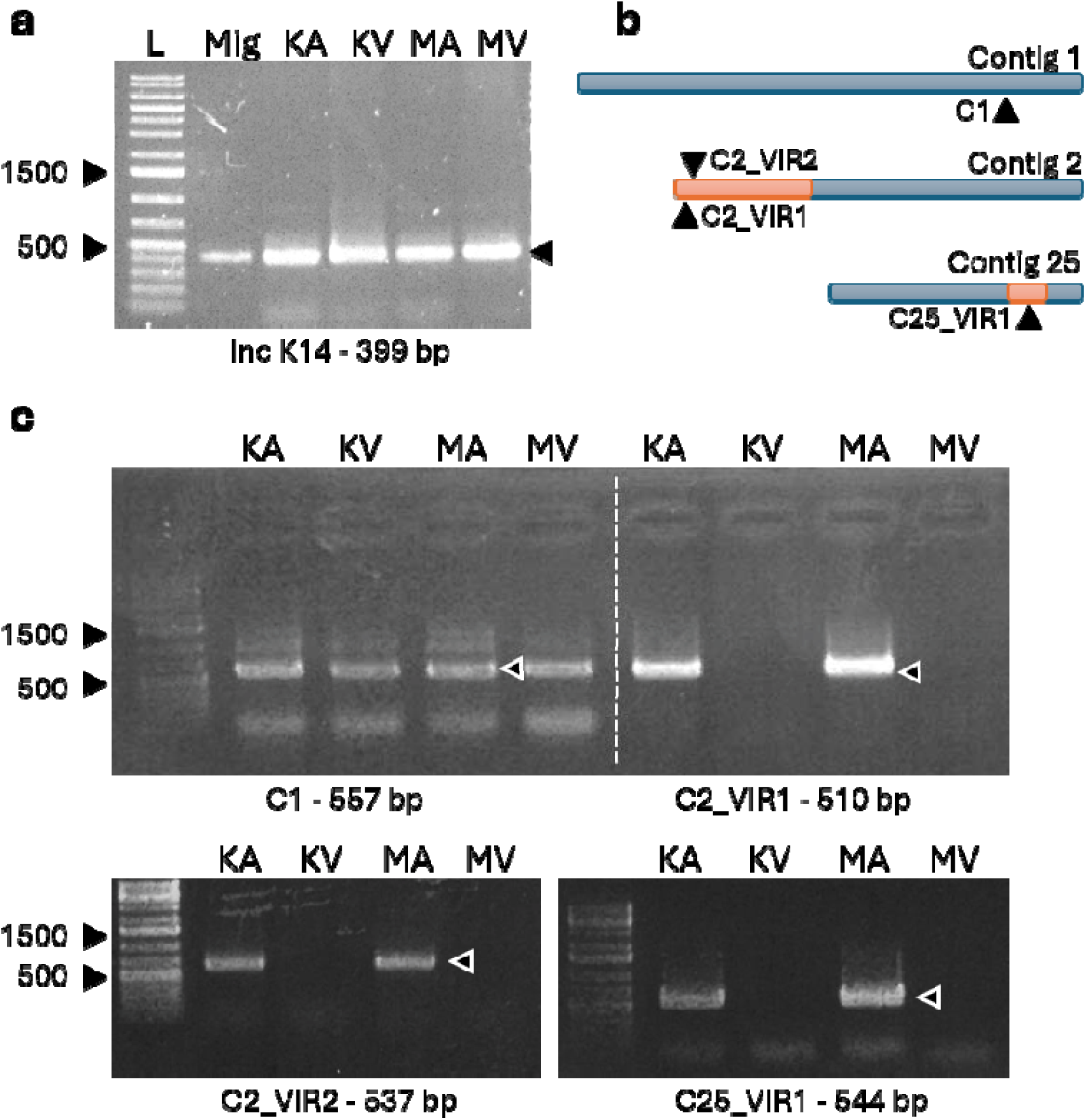
PCR validation of *M. incognita* species identity and large deletions in the two virulent isolates (KV, MV). (**a**) Species identification included a control genomic DNA of *M.incognita* (Mig) using the Inc K14 molecular marker (Randig et al., 2002). (**b**) Localizations of the molecular markers designed to validate the large deletions (orange) in contig 2 (C2_VIR1&2) and 25 (C25_VIR1) together with a control marker in a genome region without detected structural variations in contig 1 (C1). (**c**) PCR validation showing consistent bands for C1 region but an absence of band in virulent lines for C2s and C25 regions, where deletions were detected. Molecular marker names and amplicon sizes (in parenthesis) are listed below gels.

### The deleted regions of virulent isolates contain putative parasitism genes

Based on the functional annotation of the reference genome (Mota et al., 2024), lost genes in virulent isolates in contigs 2 and 25 display contrasted profiles. While 40% of the lost genes in contig 2 lack functional annotation or only secondary structure elements (e.g. coils, helices), this number rises to 80% in contig 25 **(Figure 4a)**. On contig 2, 24% of lost genes encoded various enzymes, 6% transporters, and 5% receptors, while 8% of lost genes in contig 25 encoded zinc finger proteins **(Figure 4a)**. Fifteen predicted peptidases / proteases were lost in contig 2, consistent with the GO enrichment analysis performed on DEGs. We further analyzed the distribution of genes encoding putative secreted proteins (SP-noTM), the ones with a signal peptide but with a transmembrane region (maybe secreted in vesicles, addressed to organelle membranes), transmembrane proteins (including receptors, etc) and *Meloidogyne-*specific proteins (that are not found in other genera). These are likely encoded by so-called orphan genes, hypothesized to have either highly diverged from pre-existing genes or emerged *de novo* from non-genic regions (Seçkin et al., 2025). Lost genes in contig 2 display a distribution similar to the whole predicted proteome at the exception of the transmembrane proteins which are depleted compared to the whole proteome (-5%; *□*2; P<0.05) **(Figure 4b)**. On the other hand, lost genes in contig 25 are enriched in orphan genes compared to the whole proteome, regardless of their putative *de novo* origin or not (+10%; *□*2; P<0.05) **(Figure 4b)**.

**Figure 4.**
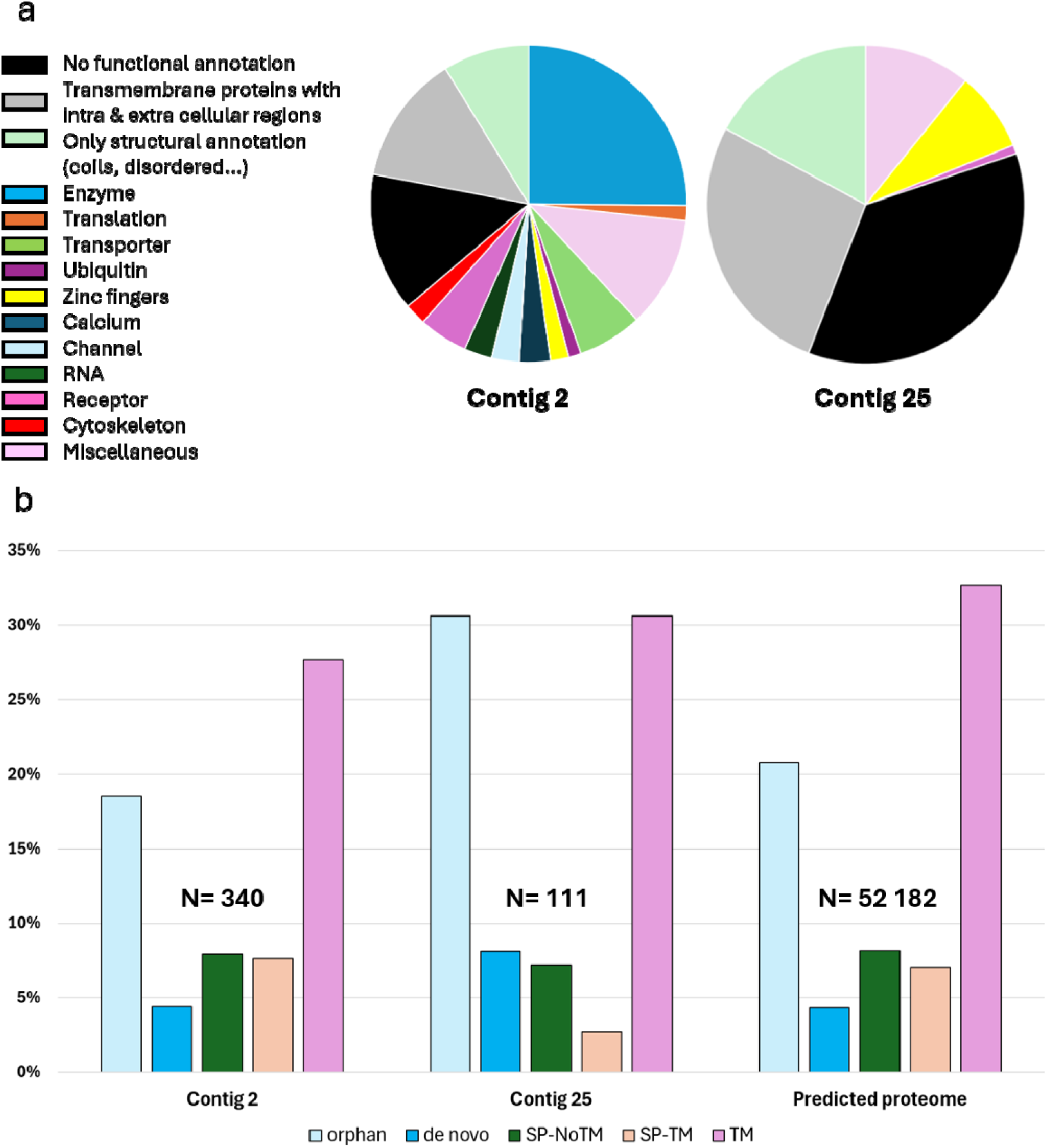
Functional annotation of the 340 and 111 lost genes in the large parallel deletions in contigs 2 and 25. (**a**) Functional annotation based on InterProScan domain identification. (**b**) Percentage of genes that are predicted to be orphans, *de novo* orphans, secreted including a signal peptide (SP) without (SPNoTM) or with (SPTM) transmembrane domain (TM).

Among lost genes, we found proteins potentially related to parasitic processes such as a chorismate mutase (Bekal et al., 2003) and about ten carbohydrate-active enzymes (CAZymes) (Danchin et al., 2010; Haegeman et al., 2011) among which some are predicted to be secreted. The full list is available in **Supplementary Table S6**. Interestingly, among the lost genes of contig 2, we identified one encoding the *M. incognita* avirulence protein MAP-1 (CAC27774, Minc_v4_shac_contig_2g0016591), which was widely studied for its putative role in parasitism (Castagnone-Sereno et al., 2009; Rosso et al., 2011; Rutter et al., 2014; Semblat et al., 2001; Tomalova et al., 2012). We functionally reannotated MAP-1 with a more recent method and database. In addition to the signal peptide in the N-terminal region, we identified a disordered proline-rich region that includes the previously described repetitive motifs (Semblat et al., 2001) and a Barwin-like endoglucanases domain, typical from expansin proteins, which mediate the loosening and extension of plant cell walls (Cosgrove, 2000). Using the long-read versions of the reference genomes of *M. incognita, M. arenaria, M. javanica*, *M. luci*, and *M. enterolobii* (Mota et al., 2024; Poullet et al., 2025), we found 3, 1, 2, 2 and 2 proteins, respectively, that strictly display this signature pattern (i.e., including a signal peptide, a disorder proline-rich region, and a Barwin-like endoglucanases domain). The polymorphism between these homologs is mainly located in the disordered proline-rich region whereas the Barwin-like endoglucanases domain is highly conserved **(Figure 5a)**. As expected, secondary and 3D structures are weakly predicted in the disordered proline-rich region but with high degree of confidence in the Barwin-like endoglucanases domain **(Figure 5b&c)**. The length of the disordered proline-rich region splits the *M. incognita* MAP homologs into three groups of sequences: (i) a short region (105 amino acid) including Minc_v4_shac_contig_2g0016581, a homolog, genetically clustered, organized in a head-to-head orientation with MAP-1, (ii) a long region (250 amino acid) including Minc_v4_shac_contig_32g0258231 and most homologs found in *Meloidogyne* species studied, and (iii) an intermediate size region (215 amino acid) including MAP-1 **(Figure 5a)**. It is worth noting that MAP-1 and its genetically linked homolog were lost in both virulent populations in our dataset. Likewise, multiple effector candidates were found in the deleted regions and were likely lost in parallel in the two virulent populations.

**Figure 5.**
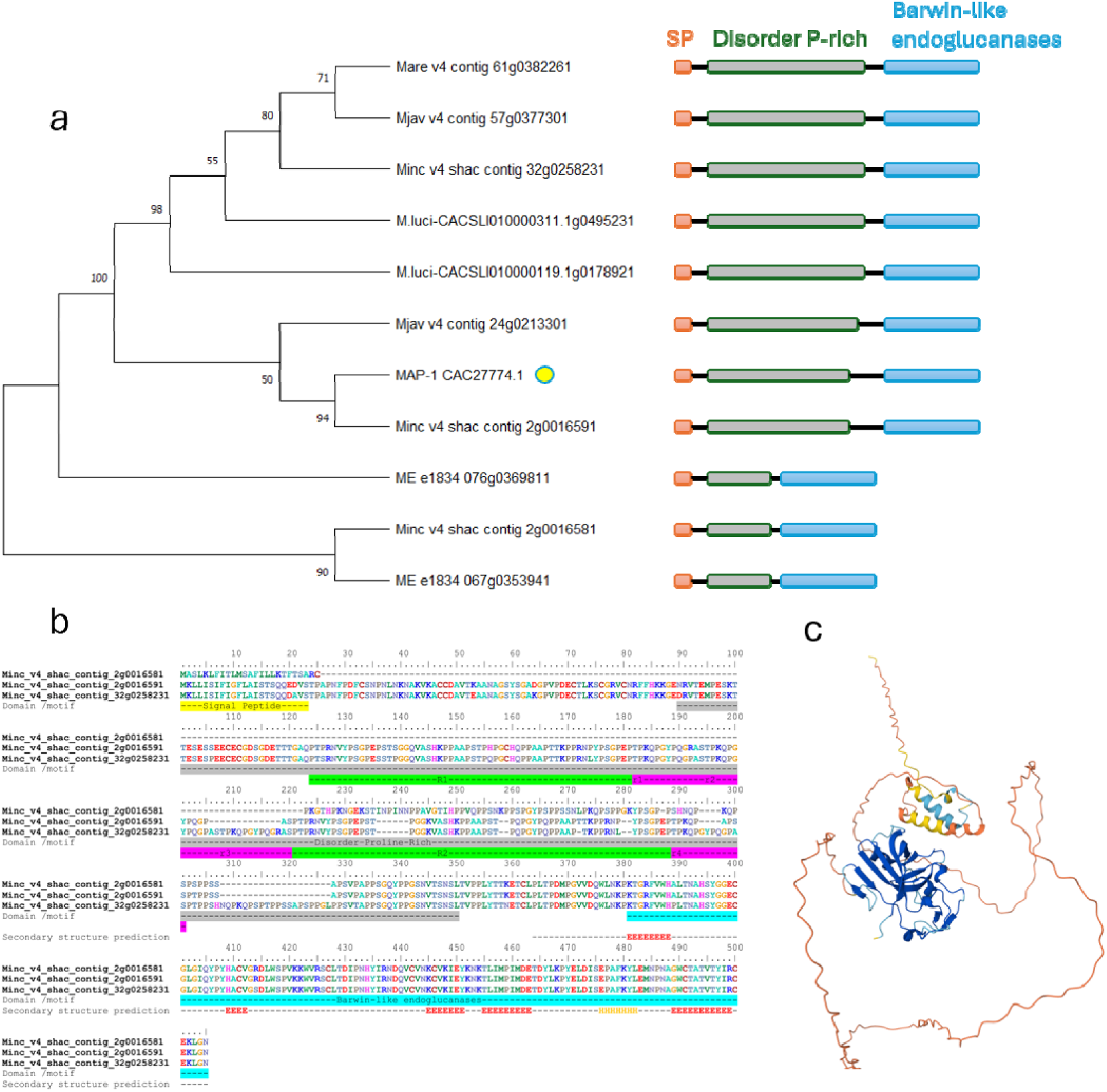
Functional and structural annotation of MAP-1 and its homologs. (**a**) Maximum likelihood phylogenetic tree of 10 homologs of MAP-1 (CAC27774.1), which is highlighted by a yellow circle, displaying the following conserved pattern: a signal peptide (SP), a variable disordered proline-rich region and a conserved Barwin like endoglucanase domain. The tree includes sequences from *M. incognita* (3), *M. arenaria* (1), *M. javanica* (2), *M. luci* (2) and *M. enterolobii* (2). (**b**) Multiple sequence alignment of MAP-1 (Minc_v4_shac_contig_2g0016591) and the two identified homologs in *M. incognita.* The alignment shows the domain / motif localization, the repeat regions in the Proline-rich region (R1, R2, r1, r2, r3 and r4) described in (Castagnone-Sereno et al., 2009), and the conserved Barwin-like endoglucanase domain. Secondary structures in the Barwin-like endoglucanase domain are predicted using JPred 4 indicating ꞵsheet (red E) and helix (orange H) structures. (**c**) 3D structure prediction of MAP-1 using Alpha fold 3. The Barwin-like endoglucanase domain, highlighted in dark blue, indicates high confidence of 3D prediction (plDDT > 90).

## DISCUSSION

The process by which root-knot nematodes acquire virulence against tomatoes bearing the *Mi*-1.2 resistance gene has remained enigmatic for decades. Studies, using experimental evolution based on avirulent/virulent near-isogenic lines of *M. incognita*, have investigated point mutations or gene copy number variation (CNV) associated with such processes (Castagnone-Sereno et al., 2019; Neveu et al., 2003; Semblat et al., 2001). Some parallel mutations, CNV, or gene expression dysregulation were previously found to be associated with virulence. Yet, the identification of massive parallel losses of genomic material in virulent isolates of distant geographic origins in the present study is a major advance towards better understanding virulence acquisition. First, it provides an explanation for the parallel gene copy losses previously observed in the same virulent isolates (Castagnone-Sereno et al., 2019; Neveu et al., 2003; Semblat et al., 2001). More importantly, this discovery suggests that virulent *M. incognita* isolates most likely escape *Mi1-*mediated recognition by plant defence systems by losing genomic regions containing hundreds of genes. Overall, the identified deletions in virulent isolates are mainly localized on two contigs, and represent *ca.* 1% of the genome and encompass 451 protein-coding genes. Therefore, virulence acquisition more likely results from loss of function than from a gain of function.

In several plant parasites and pathogens, large deletions or rearrangements affecting effector loci have been shown to enable rapid evasion of host recognition. In the poplar rust fungus *Melampsora larici-populina*, the virulence against the *RMlp7* resistance gene was driven by the combination of a point mutation and a complete deletion of 30 kb in the candidate avirulence locus (Louet et al., 2023). Similarly, in the oomycete *Plasmopara viticola*, the breakdown of the *Rpv3.1*-mediated grapevine resistance is significantly associated with genome deletions, including a 30 kb deletion encompassing potential effector-encoding genes (Paineau et al., 2024).

The ability of *M. incognita* to adapt rapidly is remarkable given its obligatory asexual reproduction. Without meiotic recombination, clonal organisms are unable to combine beneficial alleles from different individuals or populations, which theoretically limits efficacy of selection and their adaptability (Glémin et al., 2019; Neher et al., 2010). This lack of outcrossing and reduced efficiency of selection have been confirmed by previous population genomics analyses at the point mutation level (Koutsovoulos et al., 2020). Our new findings support the view that substantial genomic variations, specifically, large parallel deletions, are involved in the adaptation to resistance. However, and despite the advantage of virulent nematodes to thrive on resistant plants, they did not get the upper hand over the global root-knot nematode population in fields, overall.

These large-scale genomic losses, while beneficial in the context of escaping a resistance gene, may be counterselected in environments where susceptible or multiple hosts are available. This paradox may be explained by a reproductive fitness cost associated with virulence acquisition and the associated gene losses. Indeed, previous works have demonstrated that virulent *M. incognita* lines exhibit lower fitness on susceptible hosts compared to their avirulent counterparts (Castagnone-Sereno et al., 2007). Confirming a cost associated with acquisition of virulence, *M. incognita* virulent isolates have been shown to lose their ability to parasitize pepper, regardless of the presence or absence of the resistance gene (Djian-Caporalino et al., 2011; Kunwar et al., 2026). Interestingly, a cost associated with virulence acquisition has also been described in *M. javanica*, a closely related asexual species from the same genus (Blundell et al., 2026).

The extent of the deleted regions in *M. incognita* might be partially explained by the triploid nature of its genome. Most genes being in three copies, we hypothesize that losing a series of them could be offset by the presence of other copies. Yet, the compensation is not expected to be complete. Indeed, it has been shown that the homoeologous gene copies are not fully redundant at the expression level at different life stages or on different host plants (Blanc-Mathieu, Perfus-Barbeoch, Aury, Rocha, et al., 2017; Dai et al., 2023; Moura de Souza et al., 2026),

The discovery of parallel deletions in loci containing putative effectors across two independent lineages suggests a repeatable evolutionary trajectory towards overcoming *Mi-*1.2 resistance. This parallel evolution is a hallmark of strong selective pressure, as seen in some other global pathogens like *Zymoseptoria tritici*, where rapid adaptation to wheat host resistance is facilitated by parallel genomic changes in different production areas (H. Chen et al., 2023). The fact that the deletions fixed in the two virulent isolates encompass specific effector clusters reinforces the hypothesis that losing recognized "avirulence" factors is an effective way for pathogens to rapidly evade host immunity (Louet et al., 2023).

Among the NLR gene family, guardians of the effector-triggered immunity in plants, the *Mi-*1.2 gene is uncommon as it controls three pests (i.e. root-knot nematodes, the potato aphid, and the sweet potato whitefly) (Rossi et al., 1998). Some studies showed that the requirements for *Mi*-mediated resistance to nematodes and aphids vary (Goggin et al., 2006), raising the question whether the *Mi* protein serves as a platform to recognize multiple different effectors. More probably, *Mi*-1.2 is capable of detecting modifications on host proteins induced by effectors of different origins. The *Mi*-1.2 gene controls 13 *Meloidogyne* species, including the three most damaging and closely related species *M. incognita, M. javanica and M. arenaria*. However, it does not control the phylogenetically more distant species *M. hapla, M. fallax,* and *M. enterolobiiI.* Curiously, *M. floridensis*, which is the closest relative of *M. incognita*, is also not controlled by *Mi-*1 (Gabriel et al., 2020; Handoo et al., 2004; Marquez & Hajihassani, 2023). This suggests something present and conserved in the genomes of *M. incognita*, *M. javanica* and *M. arenaria* but absent or substantially different in *M. floridensis* might be involved in the recognition by *Mi-*1. Therefore, future comparative genomic studies based on the lost regions identified here, among these species, may contribute to unraveling the requirement for *Mi*-1.2 resistance.

Among the possible candidates, the *Meloidogyne* avirulence protein MAP-1 is particularly interesting. It was initially discovered using amplified fragment length polymorphism coupled with RT-PCR in three pairs of *M. incognita* virulent vs. avirulent near-isogenic lines (Semblat et al., 2001) The MAP-1 genomic marker was neither detected in *M. incognita* virulent lines, nor in *M. hapla*, *M. fallax*, nor nematodes from other genera, whereas amplification was observed in avirulent lines of *M. incognita*, *M. javanica* and *M. arenaria. Mi*-1-mediated resistance operates mainly via a post-penetration hypersensitive response, pointing out effectors acting and being (or their products) recognized at this specific stage. However, minor secondary effects such as localized cellular defense signals and slightly reduced juvenile attraction or penetration have also been reported (Bhattarai et al., 2007; Milligan et al., 1998; Williamson, 1998). Interestingly, the immunolocalization of MAP-1 showed a clear signal in the amphids of *M. incognita* juveniles (a secretory organ) and an accumulation in the extracellular compartment during root infection, suggesting a role in the early steps of the nematode-host interaction (Rosso et al., 2011; Semblat et al., 2001). Our functional annotation suggests that MAP-1 is a secreted expansin-like protein. Expansins mediate the loosening of plant cell walls by disrupting non-covalent bonds between cellulose microfibrils and matrix polysaccharides (Cosgrove, 2015). It is important to note that the two virulent *M. incognita* isolates lost both MAP-1 and its neighboring homologous copy. These proteins or the modifications they cause might be detected by the plant immune system. However, the virulence acquisition is unlikely to rely exclusively on MAP genes, and our annotation provides a first catalogue of candidate genes that require in-depth functional characterization to confirm their roles in the plant-nematode interaction. Understanding these molecular mechanisms is essential for the sustainable management of resistance genes and the development of innovative crop protection strategies.

Using experimental evolution assays on susceptible versus resistant host plants, we have shown large parallel genome deletions accompanying the shift from avirulence to virulence in two independent isolates - one sampled from Kursk, Russia, and the other from Morelos, Mexico. This parallel trajectory likely reflects positive selection in response to host resistance, under controlled experimental conditions. Agricultural environments are heterogeneous over space and time, with human-implemented resistance from host genes and pesticides, interacted with diverse biotic and abiotic pressure. We may therefore anticipate widespread parallel and independent emergence of genetic variations that are beyond geographic patterns. This is corroborated by recent genome-wide SNP analyses on multi-continental field populations, which shows that SNP distributions are not associated with geographic locations, attributing to multiple independent transitions and adaptations (Koutsovoulos et al., 2020). Our results thus counter common assumptions that identical genotypes share a geographic origin or that patterns inconsistent with geographic distributions are due to human-mediated translocations, while demonstrating the adaptive potential of the species facing challenging environments. Yet, should independent evolution underpin the extremely wide geographic distribution and host range of *M. incognita*, will present major challenges to both evolutionary biology and agricultural management.

These large genome deletions may represent standing genetic variation that arise before nematodes being shifted to the resistant plant, and being rapidly selected. In the molecular evolution perspective, massive gene loss could proceed through large deletions or gradual accumulation of smaller deletions (Crombez et al., 2025). Finally, these deletions may represent a source of genetic variation causing adaptative diversity (Albalat & Cañestro, 2016), or lead to mutation meltdown driving population decline (Tichkule et al., 2025). These processes and consequences of genome changes raise important evolutionary questions in regards to the species’ adaptive potential, and reiterate *M. incognita* as a model species for the investigation of adaptation of obligate asexual pathogens in rapidly changing selection pressures.

## DATA AVAILABILITY

All the raw data have been deposited in the European Nucleotide Archive under accession number: PRJEB124270. The processed data have been deposited in the publicly available French institutional repository “Recherche Data Gouv”, at this URL: https://entrepot.recherche.data.gouv.fr/dataverse/Ma-Mv-Ka-Kv/.

## ACKNOWLEDGEMENTS

We are grateful to the bioinformatics and genomics platform, BIG, Sophia Antipolis (ISC PlantBIOs, https://doi.org/10.15454/qyey-ar89) for providing help as well as computing and storage resources. We would like to thank Silvia Bottini for advice and insight in the analysis of gene expression data. This work was performed in collaboration with the GeT core facility, Toulouse, France (GeT, https://doi.org/10.15454/1.5572370921303193E12). GeT core facility was supported by France Génomique National infrastructure, funded as part of “Investissement d’avenir” program managed by Agence Nationale pour la Recherche (contract ANR-10-INBS-09).

## Authors contribution

Ana Paula Zotta Mota - APZM - Data Analysis, student supervision, manuscript writing

Hongli Yu - HY - Data analysis, figures generation, manuscript revision

Maxime Multari -MM - Transcriptomic analysis and interpretation

Rahim Hassanaly Goulamhoussen - RHG - Biological material production, DNA / RNA extraction, conceived part of the experiments, manuscript revision

Laetitia Perfus-Barbeoch - LPB - Conceptualization, funding acquisition, biological material production, DNA / RNA extraction, manuscript revision

Sophie Valière - AVITI library preparation and sequencing

Kory Jason - KJ - AVITI genome data processing and QC

Philippe Castagnone-Sereno - PC-S - funding acquisition

Pierre Abad - PA - Funding acquisition, conceptualization, manuscript revision

Cyril Van Ghelder - CVG - Molecular validation, manuscript writing

Etienne G J Danchin - EGJD - Project coordination / idea, conceptualization, results interpretation, supervision of students and postdocs, manuscript revision, funding acquisition

