## Supplementary Figure 1 for "Large parallel deletions are associated with massive gene losses and virulence acquisition in *Meloidogyne incognita*"

**Supplementary Data**

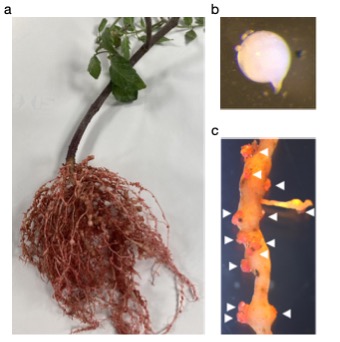

**Figure S1.** Symptoms observed on *Mi-1.2-*carrying tomatoes (*S. lycopersicum* cv. Piersol) caused by the virulent population of *M. incognita* MV (Morelos virulent). (**a**) The root system is covered by galls. (**b**) a female isolated from the root system. (**c**) Infected root magnification showing egg masses (white arrows) stained with eosine.

**Figures S2–5** are submitted as a separate file.

**Tables S2–6** are submitted as a separate file.
