## Supplementary Figures S2-S5 for "Large parallel deletions are associated with massive gene losses and virulence acquisition in *Meloidogyne incognita*"

Figure S2. Mapping depth of reads on contig 1–100 (x-axis; position is in Mb) for avirulent KA (blue) and virulent KV (red) populations. Reads were sequenced from Nanopore (NA). Depth (y-axis) is the averaged mapping depth of reads per kb window and per Gb bam file size.

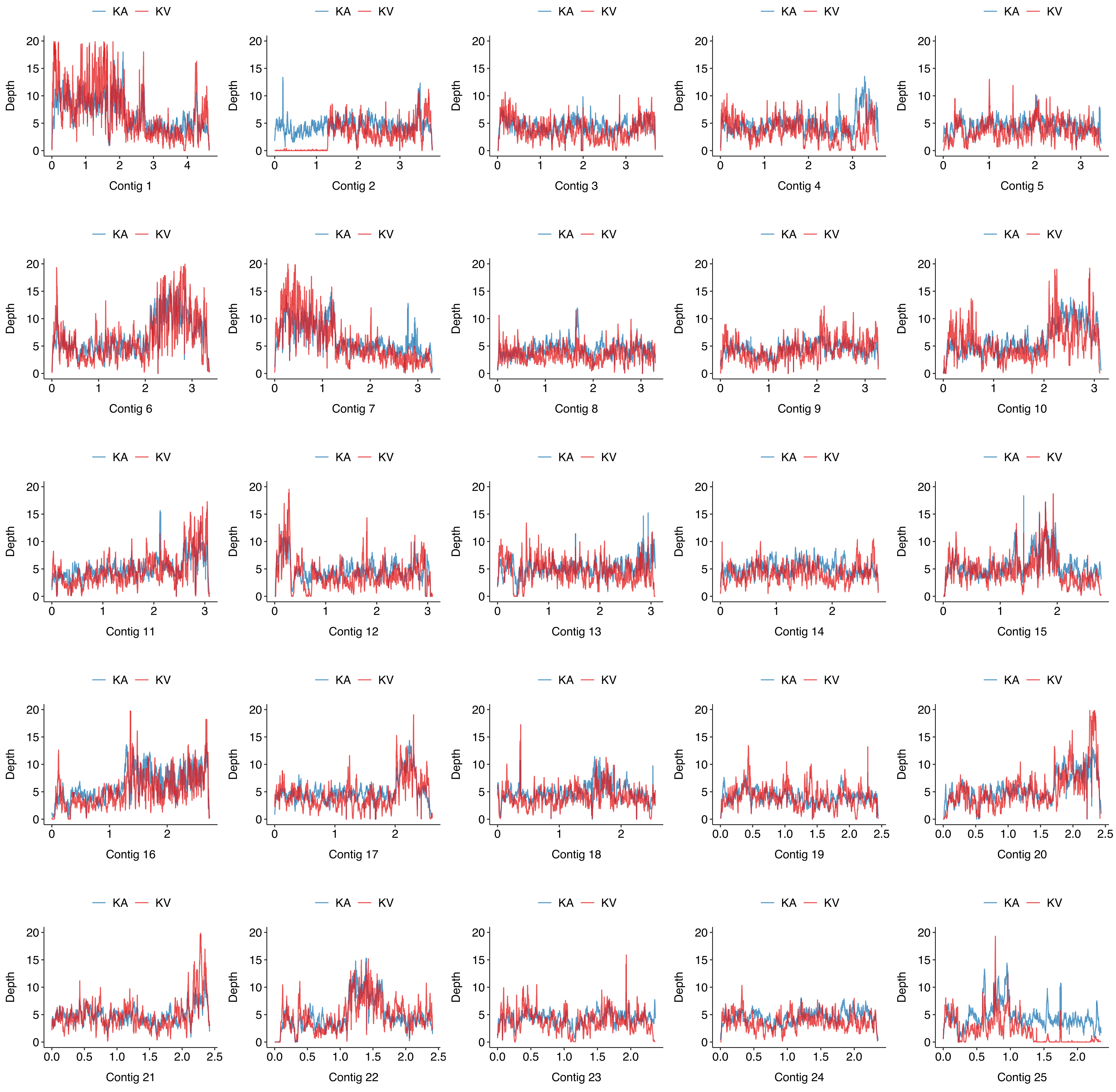

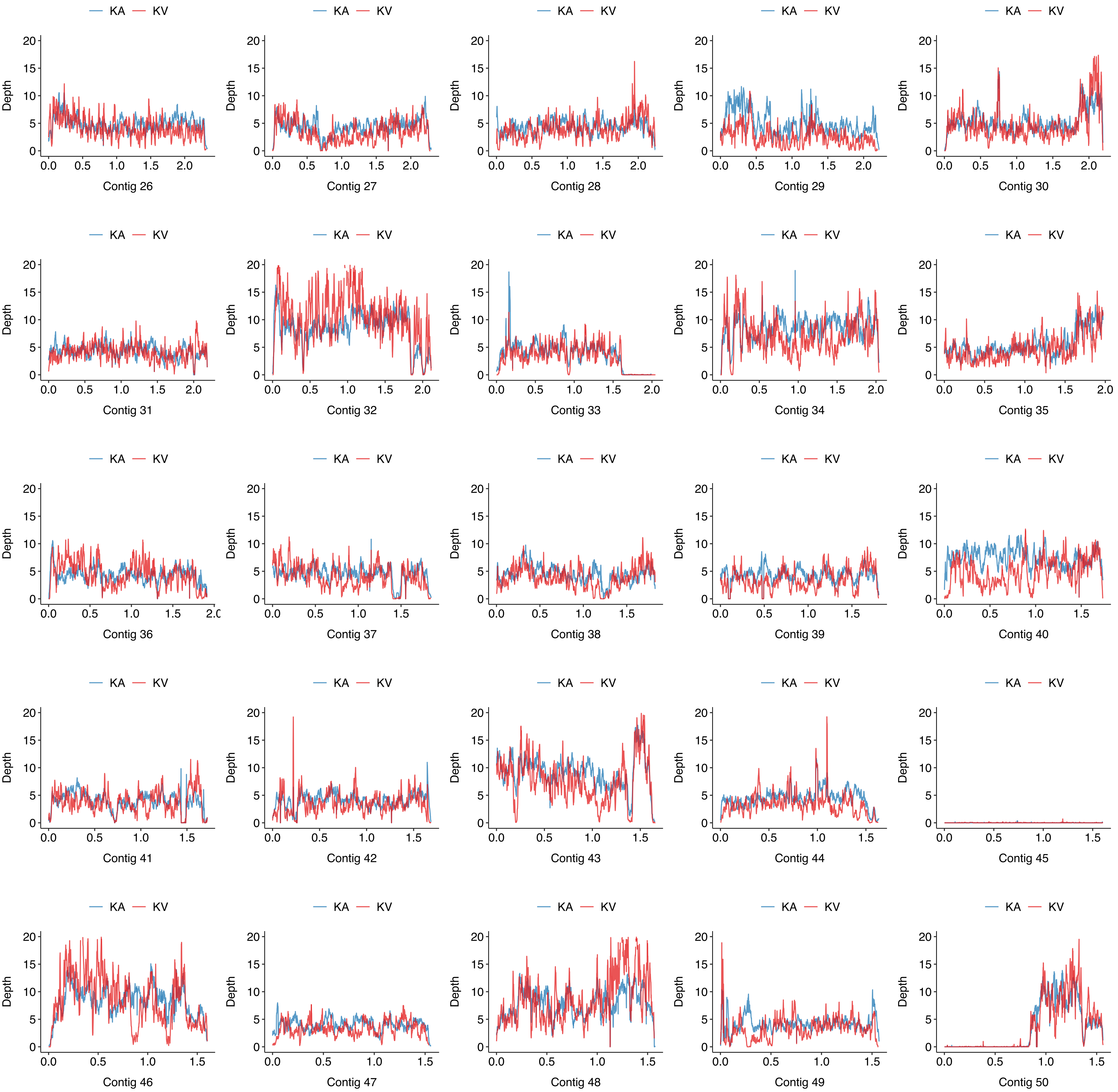

KA
KV
Depth
Contig 26
Contig 27
Contig 28
Contig 29
Contig 30
Contig 31
Contig 32
Contig 33
Contig 34
Contig 35
Contig 36
Contig 37
Contig 38
Contig 39
Contig 40
Contig 41
Contig 42
Contig 43
Contig 44
Contig 45
Contig 46
Contig 47
Contig 48
Contig 49
Contig 50

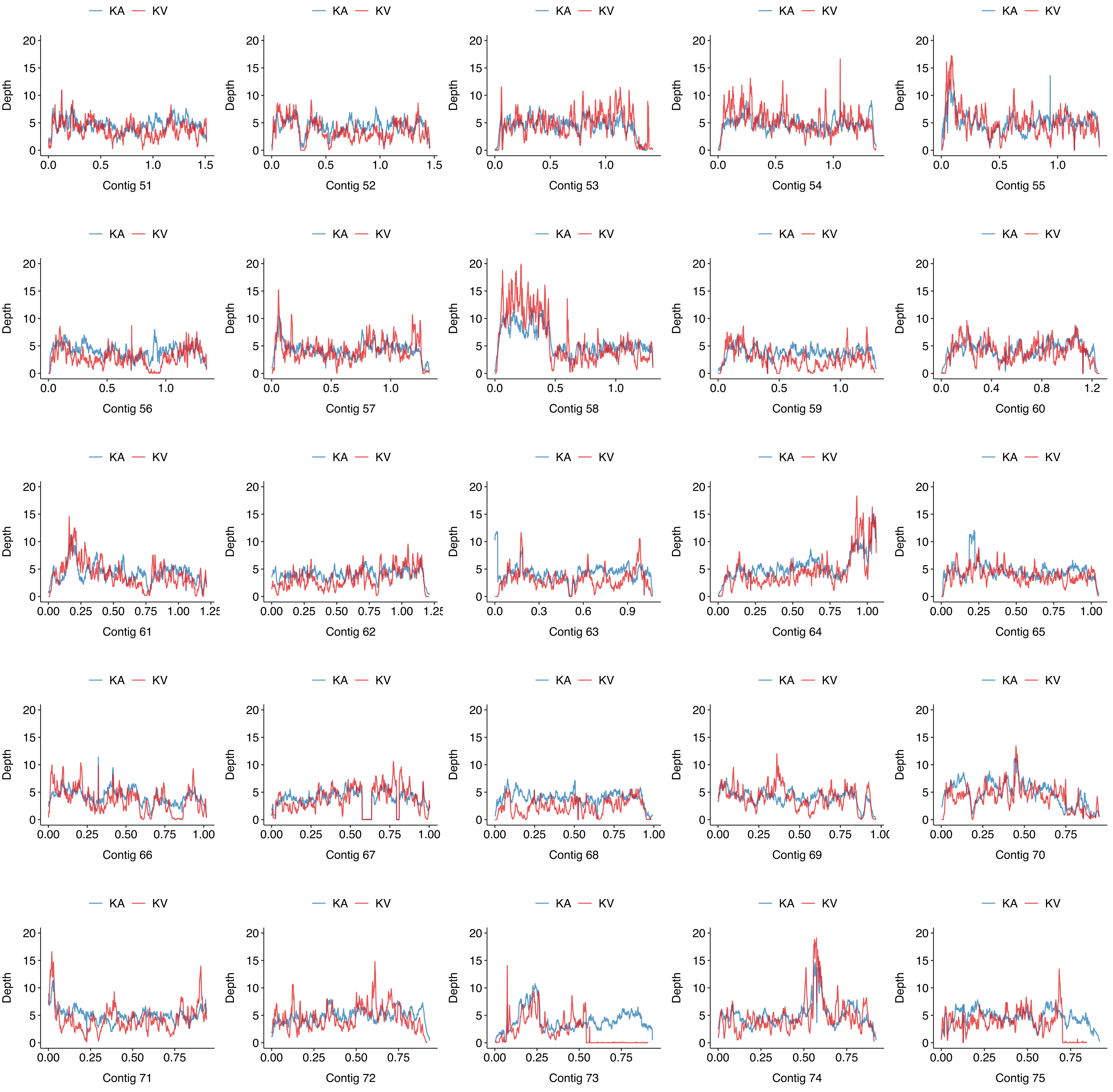

KA
KV
Depth
Contig 51
Contig 52
Contig 53
Contig 54
Contig 55
Contig 56
Contig 57
Contig 58
Contig 59
Contig 60
Contig 61
Contig 62
Contig 63
Contig 64
Contig 65
Contig 66
Contig 67
Contig 68
Contig 69
Contig 70
Contig 71
Contig 72
Contig 73
Contig 74
Contig 75

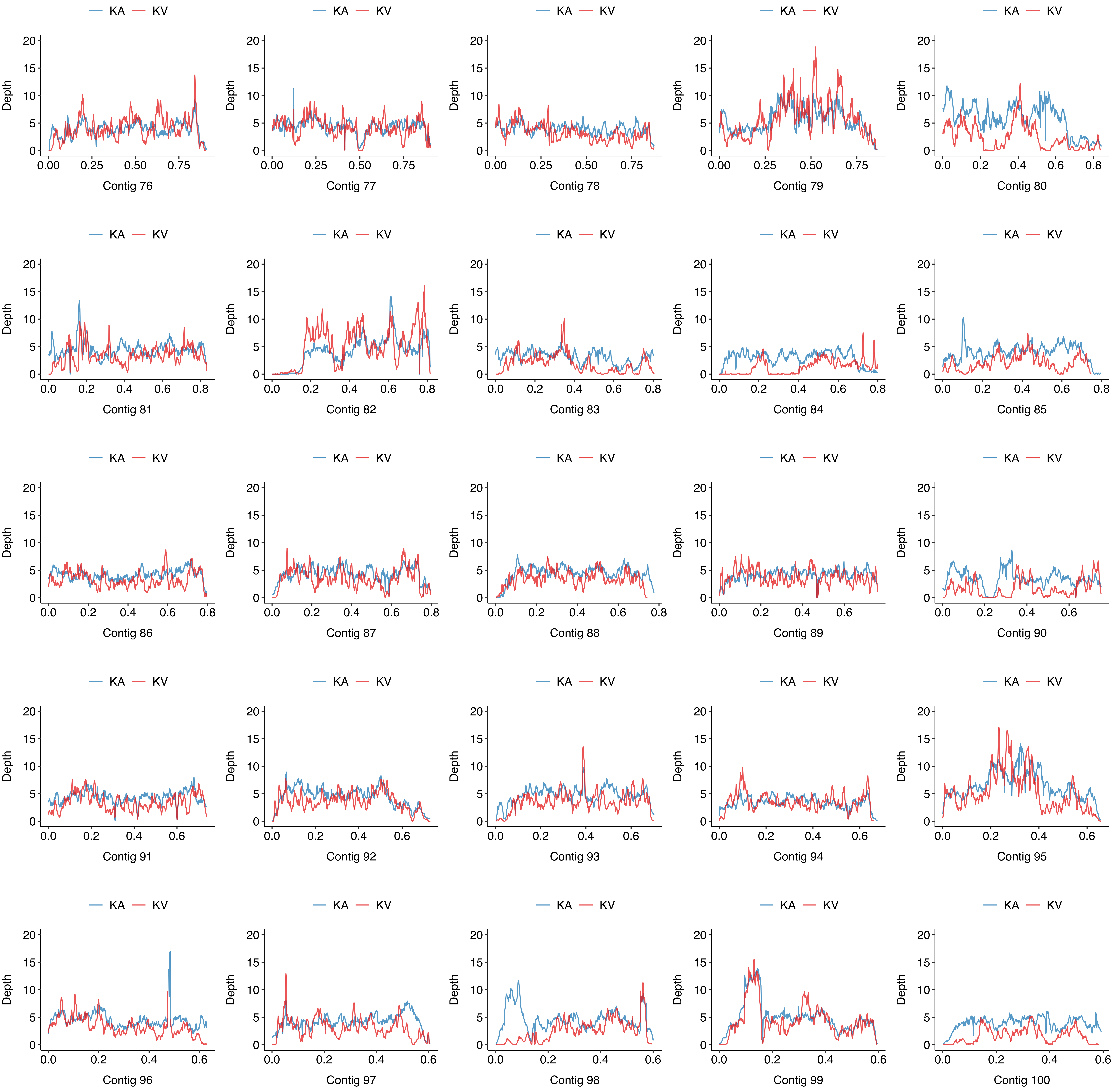

KA
KV
Depth
20
15
10
5
0
0.00
0.25
0.50
0.75
0.0
0.2
0.4
0.6
0.8
Contig 76
Contig 77
Contig 78
Contig 79
Contig 80
Contig 81
Contig 82
Contig 83
Contig 84
Contig 85
Contig 86
Contig 87
Contig 88
Contig 89
Contig 90
Contig 91
Contig 92
Contig 93
Contig 94
Contig 95
Contig 96
Contig 97
Contig 98
Contig 99
Contig 100

Figure S3. Mapping depth of reads on contig 1–100 (x-axis; position is in Mb) for avirulent KA (blue) and virulent KV (red) populations. Reads were sequenced from AVITI (AV). Depth (y-axis) is the averaged mapping depth of reads per kb window and per Gb bam file size.

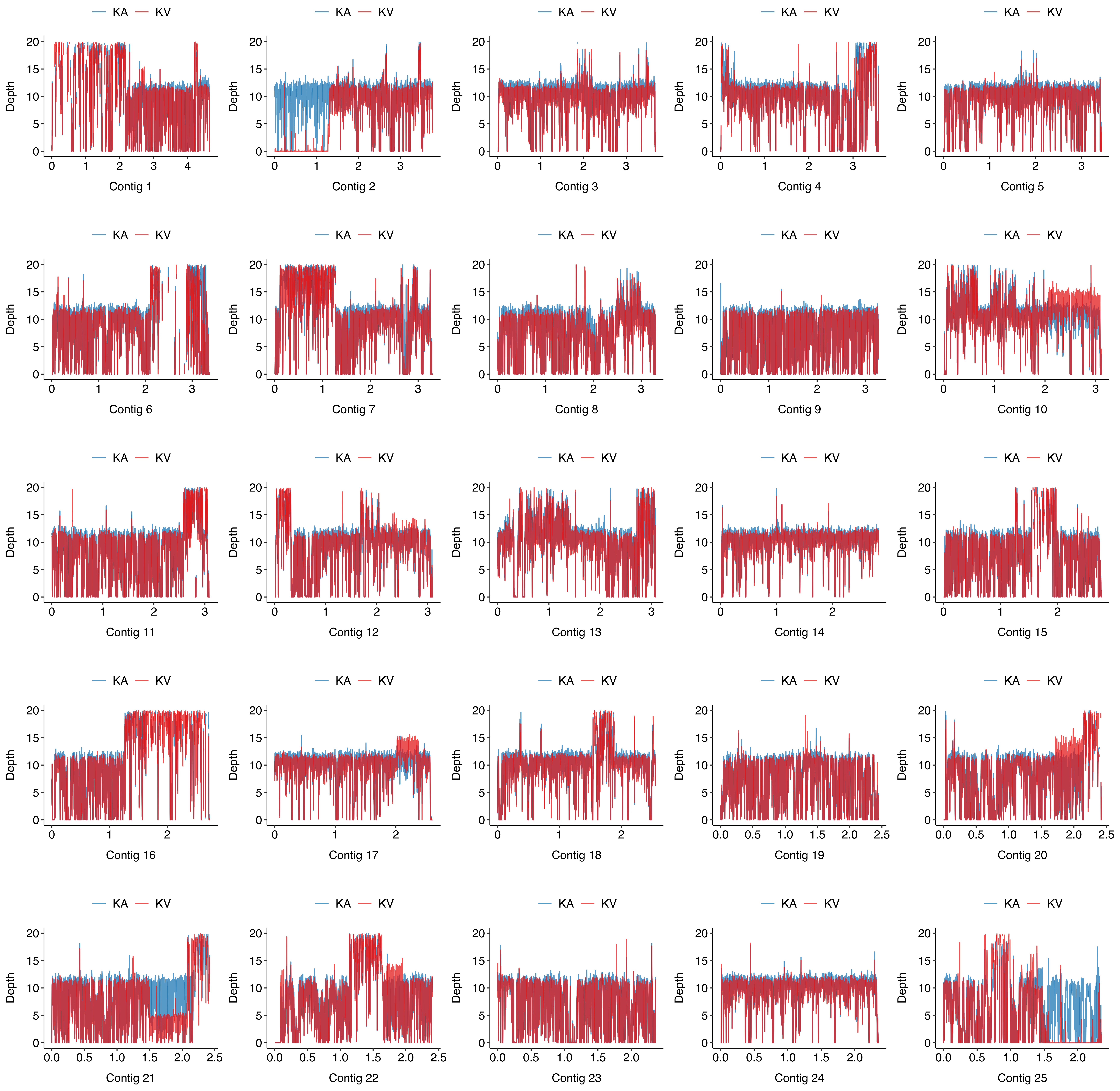

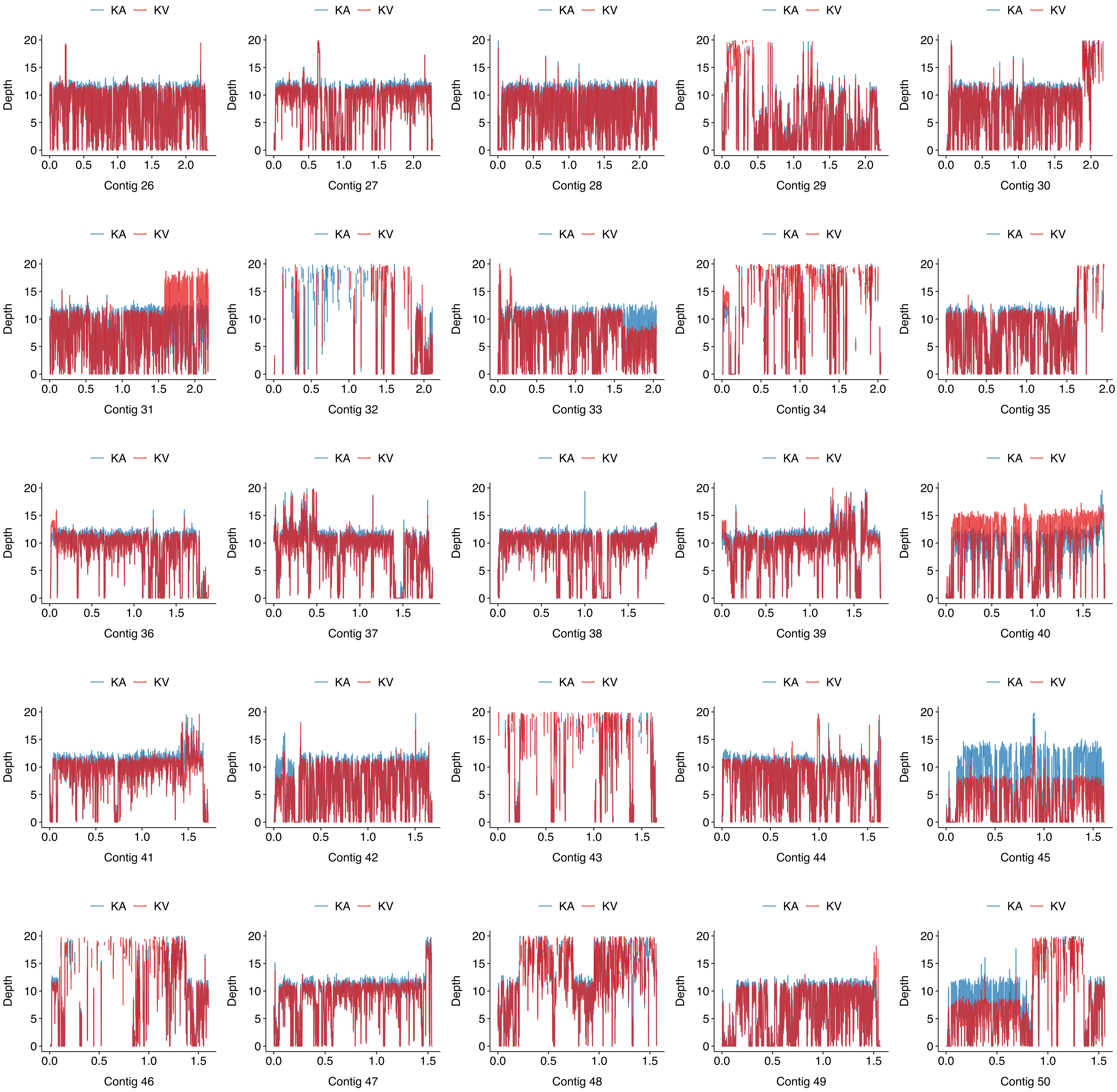
KA
KV
Depth
Contig 26
Contig 27
Contig 28
Contig 29
Contig 30
Contig 31
Contig 32
Contig 33
Contig 34
Contig 35
Contig 36
Contig 37
Contig 38
Contig 39
Contig 40
Contig 41
Contig 42
Contig 43
Contig 44
Contig 45
Contig 46
Contig 47
Contig 48
Contig 49
Contig 50

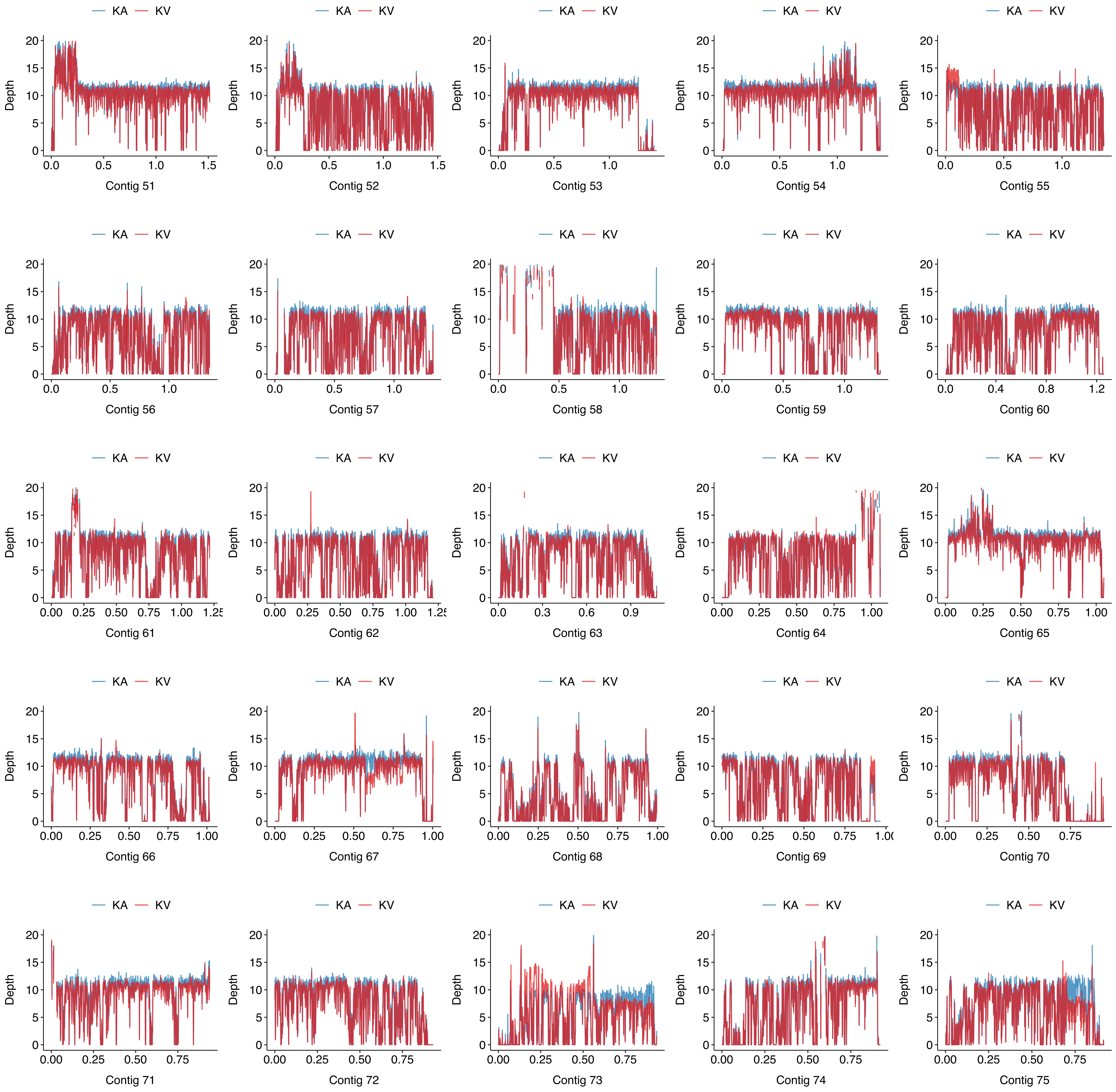

KA
KV
Depth
Contig 51
Contig 52
Contig 53
Contig 54
Contig 55
Contig 56
Contig 57
Contig 58
Contig 59
Contig 60
Contig 61
Contig 62
Contig 63
Contig 64
Contig 65
Contig 66
Contig 67
Contig 68
Contig 69
Contig 70
Contig 71
Contig 72
Contig 73
Contig 74
Contig 75

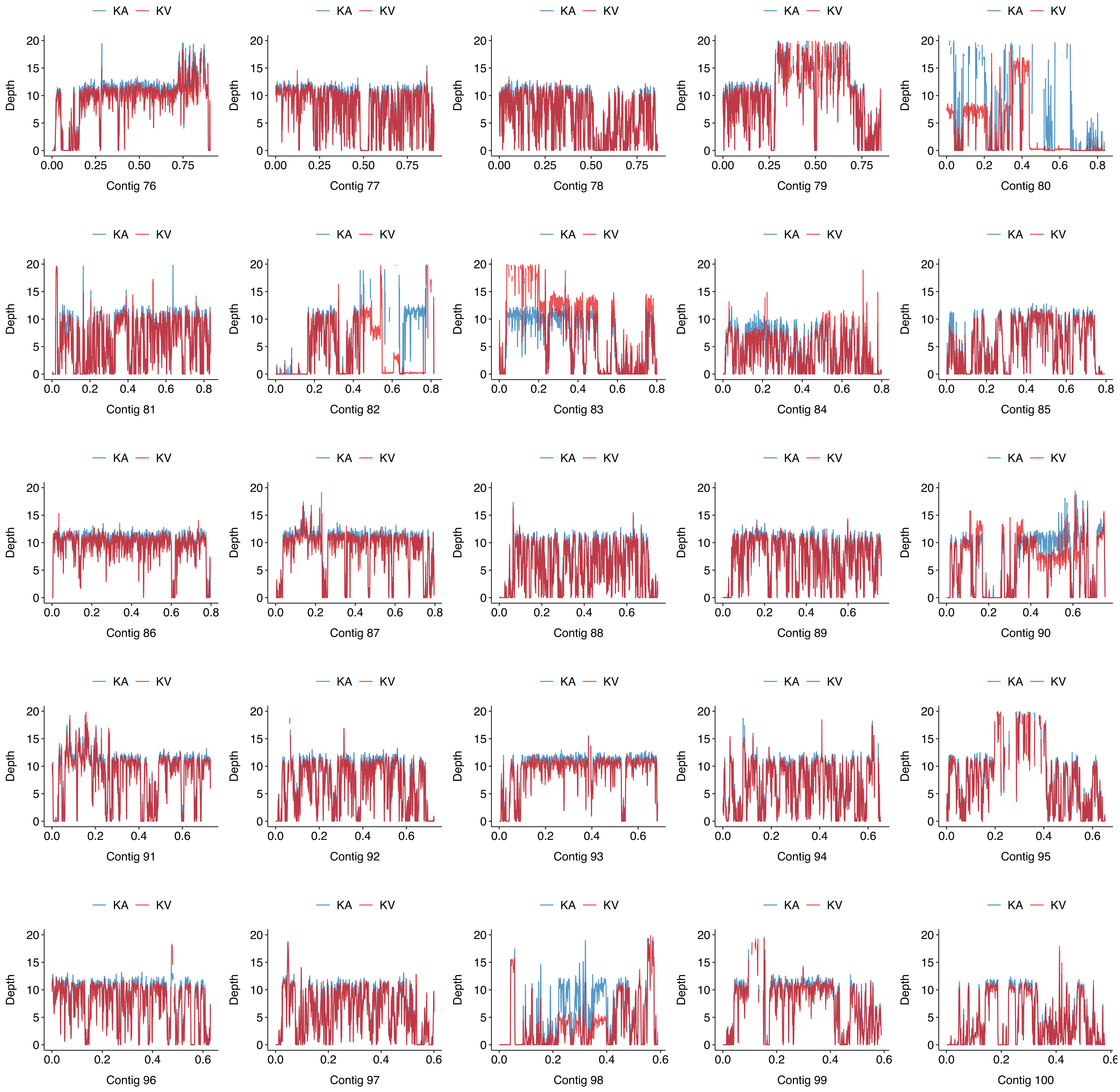
KA
KV
Depth
Contig 76
Contig 77
Contig 78
Contig 79
Contig 80
Contig 81
Contig 82
Contig 83
Contig 84
Contig 85
Contig 86
Contig 87
Contig 88
Contig 89
Contig 90
Contig 91
Contig 92
Contig 93
Contig 94
Contig 95
Contig 96
Contig 97
Contig 98
Contig 99
Contig 100

Figure S4. Mapping depth of reads on contig 1–100 (x-axis; position is in Mb) for avirulent MA (blue) and virulent MV (red) populations. Reads were sequenced from Nanopore (NA). Depth (y-axis) is the averaged mapping depth of reads per kb window and per Gb bam file size.

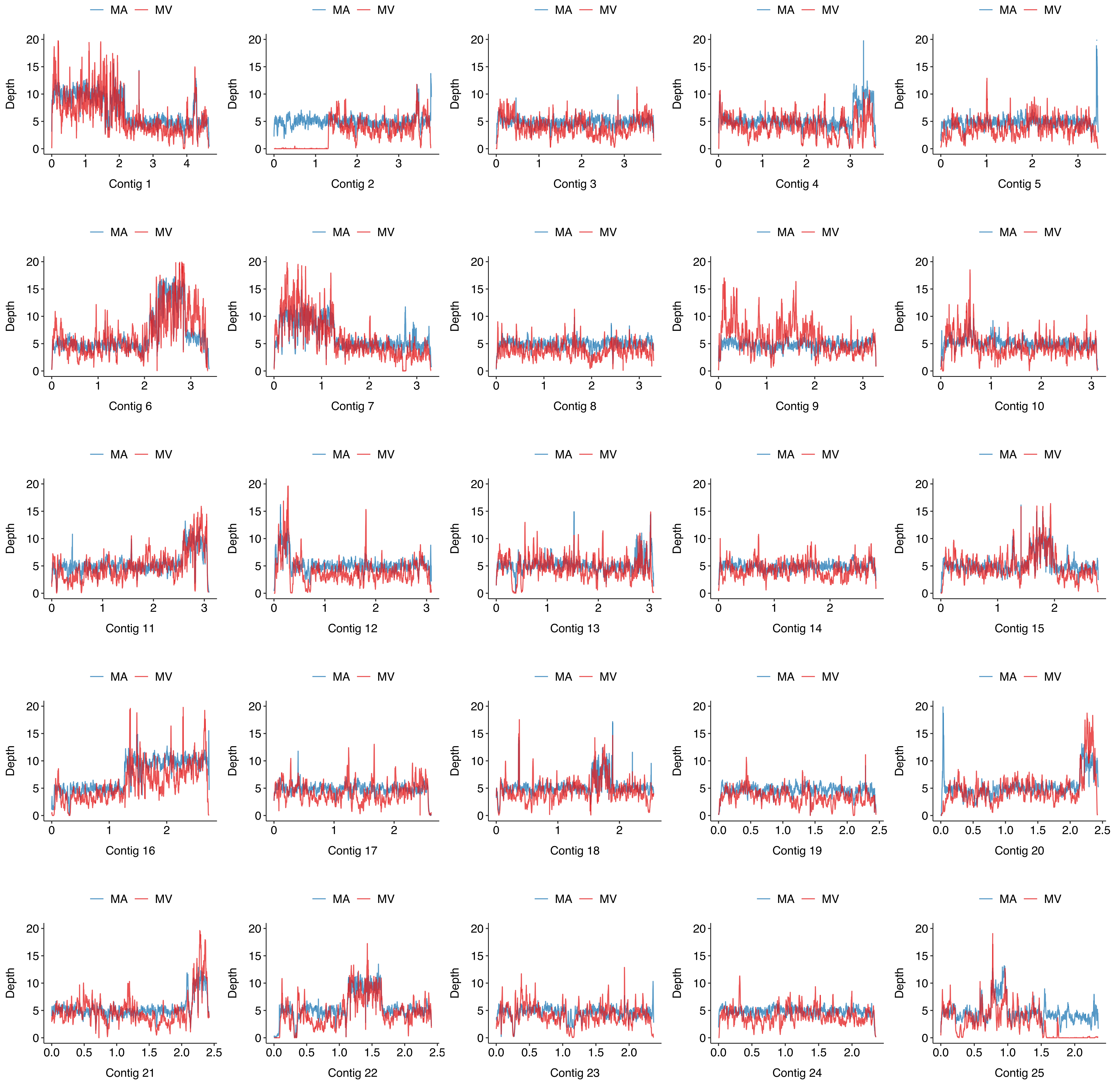

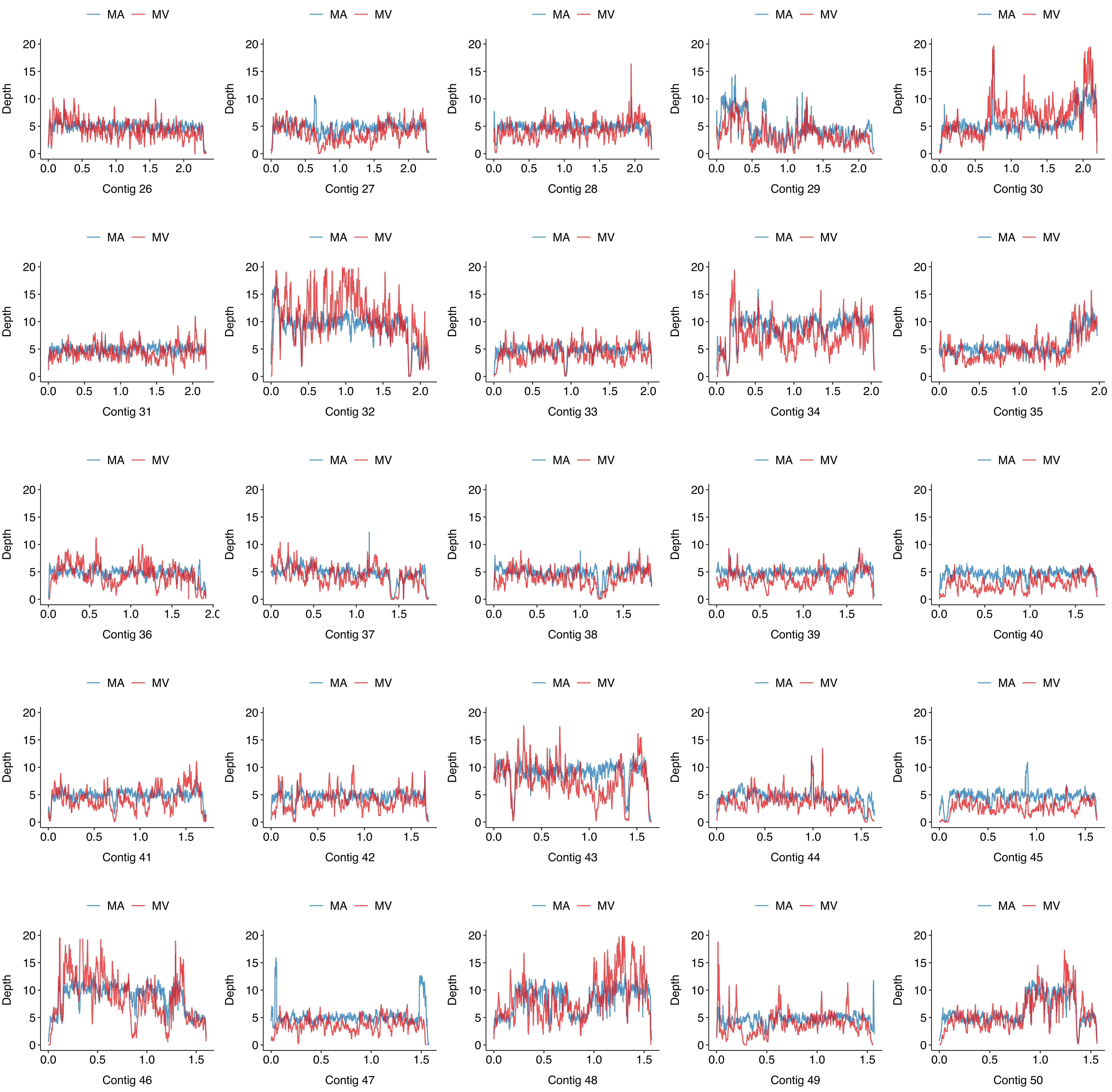

MA
MV
Depth
0
5
10
15
20
0.0
0.5
1.0
1.5
2.0
Contig 26
Contig 27
Contig 28
Contig 29
Contig 30
Contig 31
Contig 32
Contig 33
Contig 34
Contig 35
Contig 36
Contig 37
Contig 38
Contig 39
Contig 40
Contig 41
Contig 42
Contig 43
Contig 44
Contig 45
Contig 46
Contig 47
Contig 48
Contig 49
Contig 50

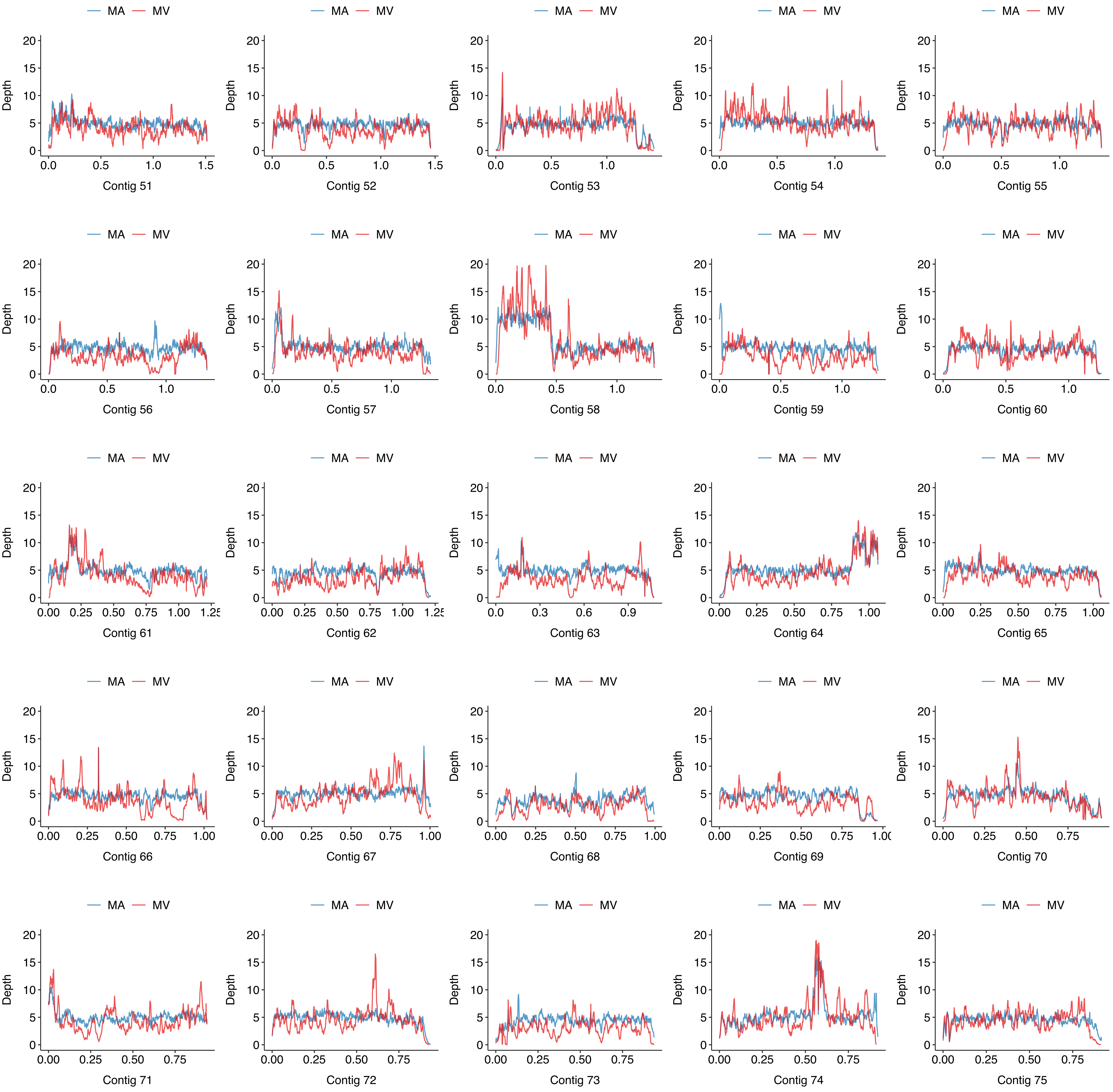
MA MV
Depth
Contig 51
Contig 52
Contig 53
Contig 54
Contig 55
Contig 56
Contig 57
Contig 58
Contig 59
Contig 60
Contig 61
Contig 62
Contig 63
Contig 64
Contig 65
Contig 66
Contig 67
Contig 68
Contig 69
Contig 70
Contig 71
Contig 72
Contig 73
Contig 74
Contig 75

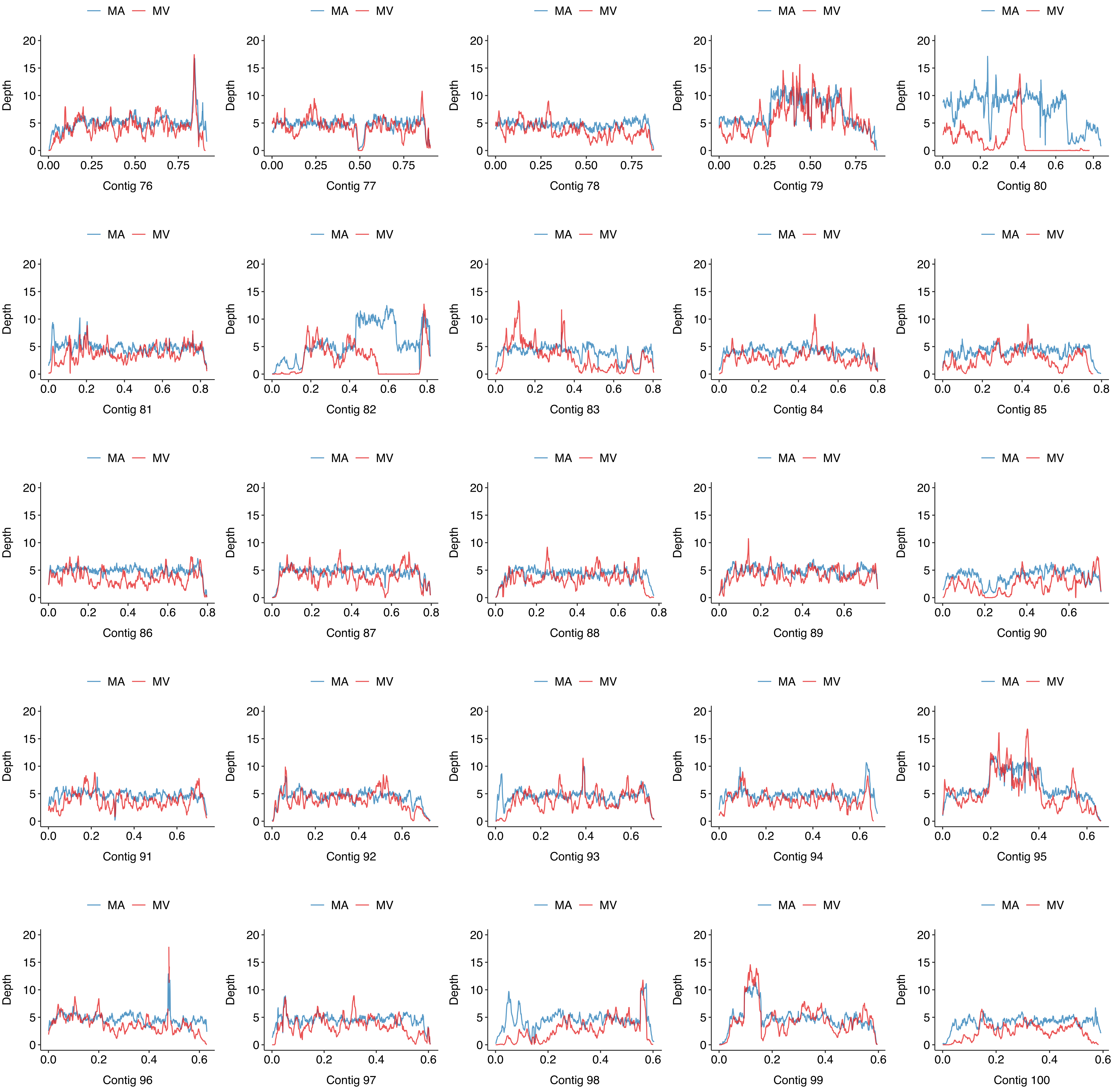
MA
MV
Depth
Contig 76
Contig 77
Contig 78
Contig 79
Contig 80
Contig 81
Contig 82
Contig 83
Contig 84
Contig 85
Contig 86
Contig 87
Contig 88
Contig 89
Contig 90
Contig 91
Contig 92
Contig 93
Contig 94
Contig 95
Contig 96
Contig 97
Contig 98
Contig 99
Contig 100

Figure S5. Mapping depth of reads on contig 1–100 (x-axis; position is in Mb) for avirulent MA (blue) and virulent MV (red) populations. Reads were sequenced from AVITI (AV). Depth (y-axis) is the averaged mapping depth of reads per kb window and per Gb bam file size.

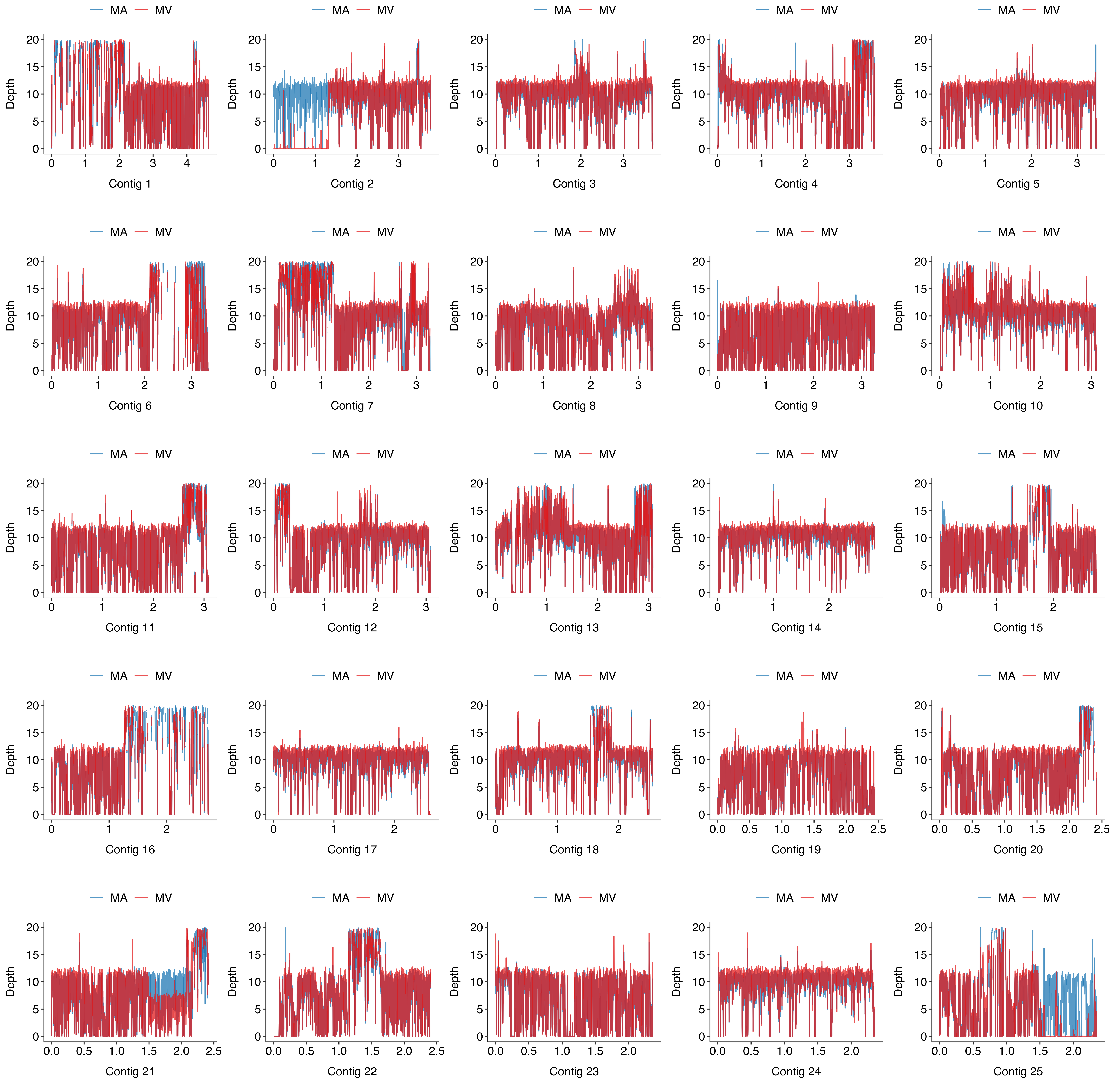

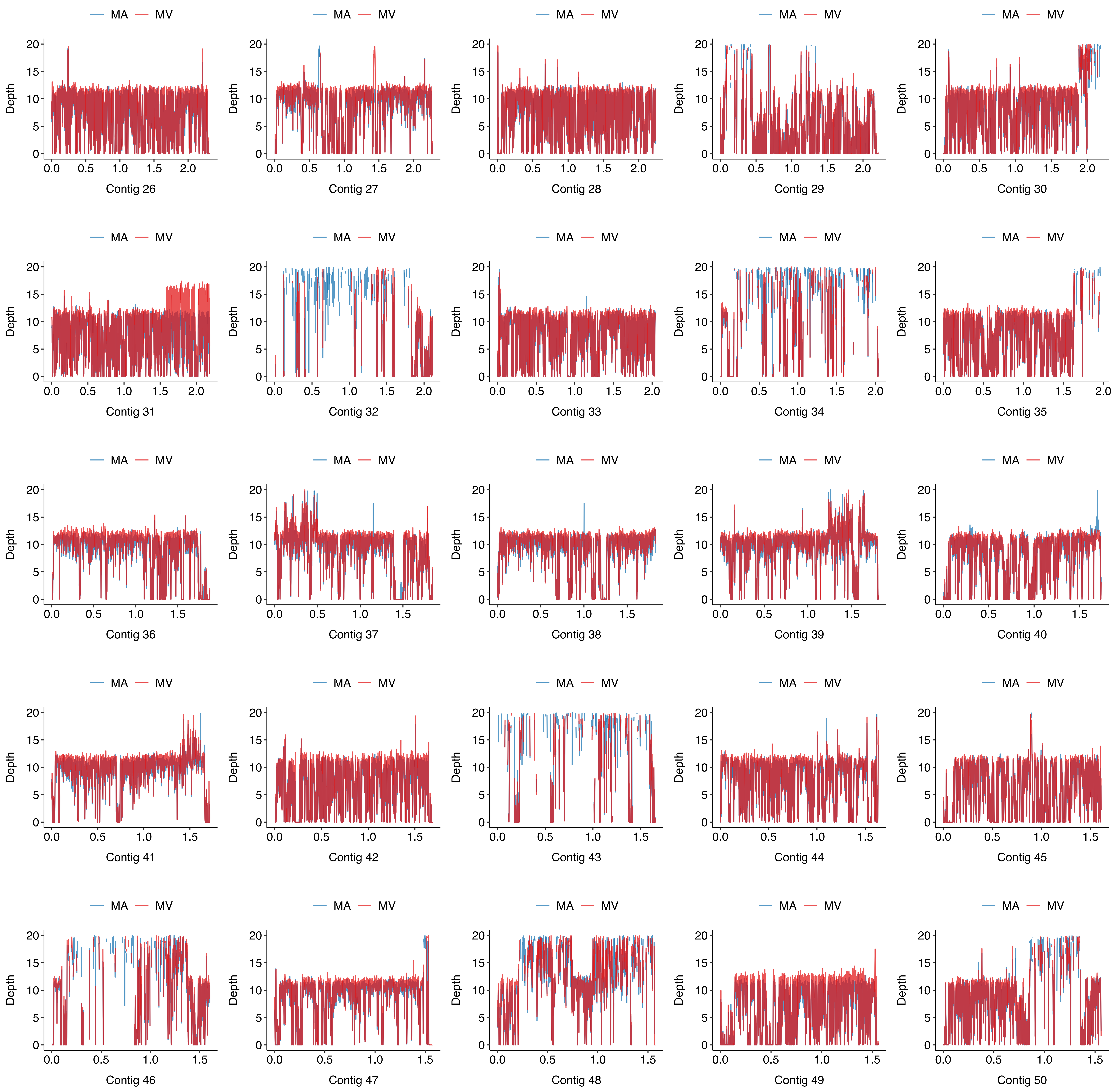
MA
MV
Depth
Contig 26
Contig 27
Contig 28
Contig 29
Contig 30
Contig 31
Contig 32
Contig 33
Contig 34
Contig 35
Contig 36
Contig 37
Contig 38
Contig 39
Contig 40
Contig 41
Contig 42
Contig 43
Contig 44
Contig 45
Contig 46
Contig 47
Contig 48
Contig 49
Contig 50

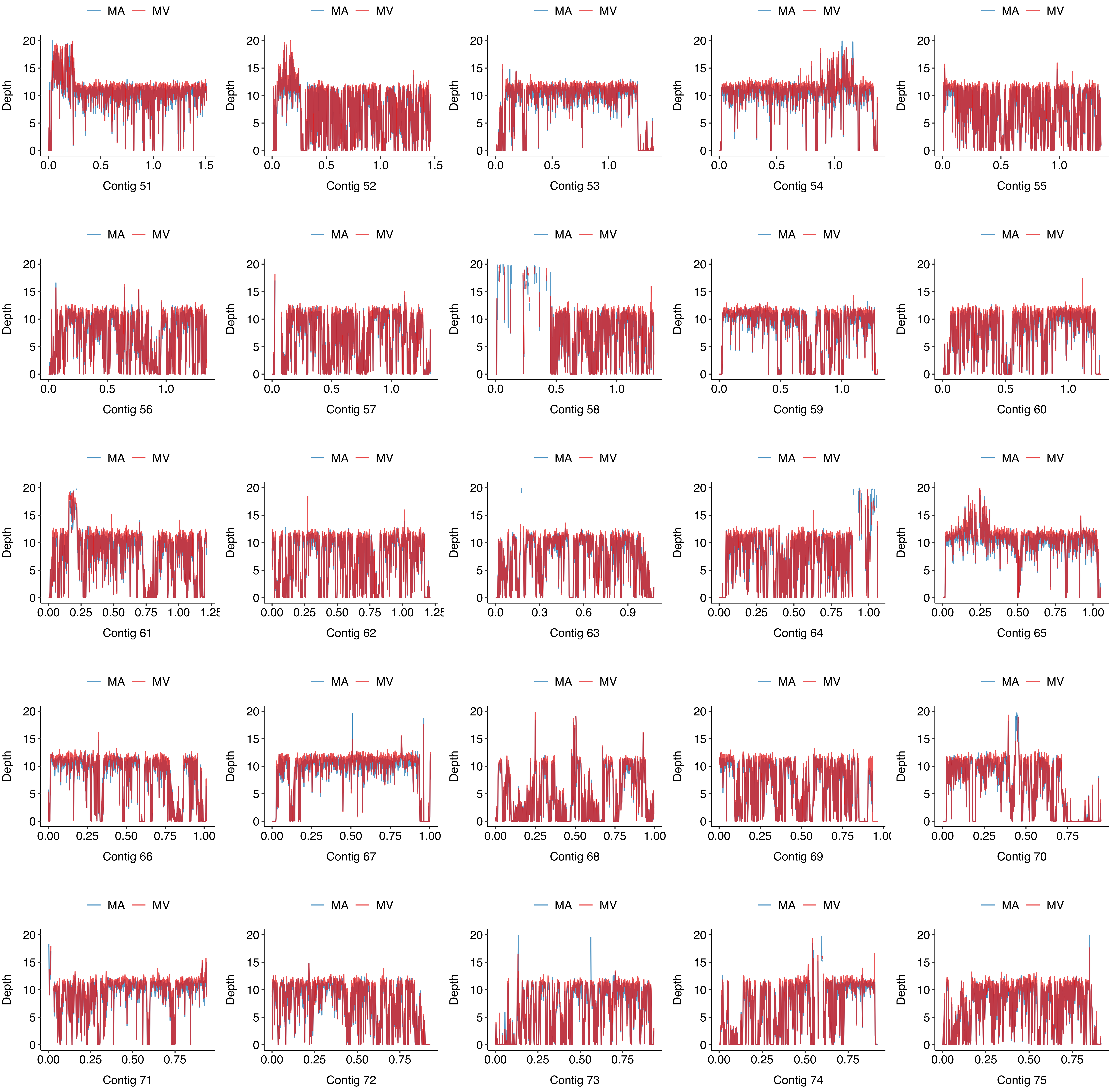

MA
MV
Depth
Contig 51
Contig 52
Contig 53
Contig 54
Contig 55
Contig 56
Contig 57
Contig 58
Contig 59
Contig 60
Contig 61
Contig 62
Contig 63
Contig 64
Contig 65
Contig 66
Contig 67
Contig 68
Contig 69
Contig 70
Contig 71
Contig 72
Contig 73
Contig 74
Contig 75

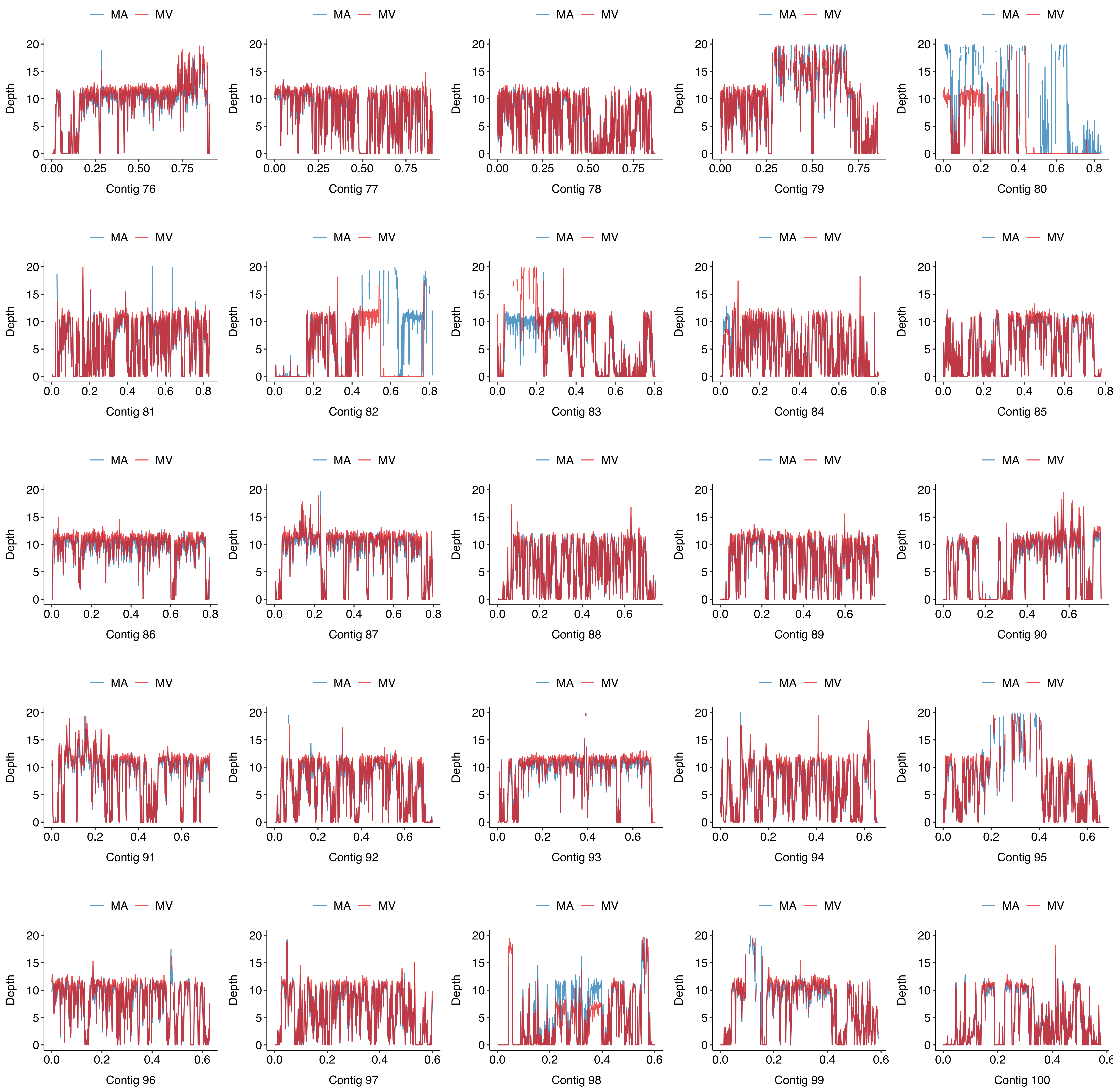
MA
MV
Depth
Contig 76
Contig 77
Contig 78
Contig 79
Contig 80
Contig 81
Contig 82
Contig 83
Contig 84
Contig 85
Contig 86
Contig 87
Contig 88
Contig 89
Contig 90
Contig 91
Contig 92
Contig 93
Contig 94
Contig 95
Contig 96
Contig 97
Contig 98
Contig 99
Contig 100
